# Gut microbiome disturbances of altricial Blue and Great tit nestlings are countered by continuous microbial inoculations from parental microbiomes

**DOI:** 10.1101/2022.02.20.481211

**Authors:** David Diez-Méndez, Kasun H. Bodawatta, Inga Freiberga, Irena Klečková, Knud A. Jønsson, Michael Poulsen, Katerina Sam

**Author notes:** These authors contributed equally to this work (David Diez-Méndez, Kasun H. Bodawatta).

## Abstract

Gut microbial communities are complex and heterogeneous and play critical roles for animal hosts. Early-life disruptions to microbiome establishment can negatively impact host fitness and development. However, the consequences of such early-life disruptions are unknown in wild birds. To help fill this gap, after validating the disruptive influence of antibiotic and probiotic treatments on the gut microbiome in adult Great tits (*Parus major*) (efficacy experiment), we investigated the effect of continuous early-life gut microbiome disruptions on the establishment and development of gut communities in wild Great and Blue tit (*Cyanistes caeruleus*) nestlings (field experiment). Despite negative impacts of treatments on microbial alpha and beta diversities in the efficacy experiment, treatment did not affect the composition of nestling microbiomes in the field experiment. Independent of treatment, nestling gut microbiomes of both species grouped by brood, sharing high numbers of bacterial taxa with both the nest environment and their mother. The distance between nests increased inter-brood microbiome dissimilarity, but only in Great tits, indicating species-specific influence of environment on microbiomes. The strong maternal effect, driven by continuous recolonization from the nest environment and vertical transfer of microbes during feeding thus appear to provide resilience towards early-life disruptions in nestling gut microbiomes.

## Introduction

Complex and heterogeneous gut microbial communities affect vertebrate host physiology, development and behavior, with ramifications for host ecology and evolution [1–5]. Early-life establishment of a functioning consortium of gut symbionts is critical for microbiome structure and function later in life [6–8]. Consequently, disruptions to early-life microbiome assembly processes can negatively impact host fitness and health by altering immune system development, increasing the probability of autoimmune diseases, and reducing resistance to parasitic infections [3, 9–12]. Oviparous birds (class Aves) acquire their initial gut symbionts after hatching, although the sterile nature of eggs and the transfer of maternal microbiota during egg formation is still controversial [13]. Post-hatching, parental and nest microbiomes [14, 15] along with diet [7, 16–18] and habitat [19–23] are thought to be the major factors shaping avian gut microbiomes.

The drivers and trajectory of establishment of the avian gut microbiomes in early life could rely on developmental patterns, from precocial to altricial species. Precocial chicks leave the nest area and feed independently shortly after hatching, whereas altricial chicks spend the brooding period within the nest and are fed directly by the parents [24]. Thus, the gut microbiome establishment of precocial chicks tends to be strongly influenced by the feeding environment, resulting in similar microbiome structures within and between broods [25]. In contrast, gut microbiomes of altricial species tend to be more similar within than between broods [26–28], possibly influenced by mutually non-exclusive vertical transmission of bacteria from parents during feeding events [15, 17] and environmental transfer of microbiomes from food items and the nest [6, 29, 30].

Gut microbiome disruptions have been carried out for decades in the poultry industry by applying antibiotics [31, 32]. In recent years, probiotics [33] have been used as growth promoters, where treatments lead to an increase in weight gain and feed efficiency [34]. Comparisons of treatment outcomes between poultry and wild birds are difficult, as living environments, diets and microbial communities of chickens have been selected by humans for decades [5]. Nevertheless, the limited number of studies that have investigated the effect of antibiotic treatments on wild chick microbiomes have demonstrated an increase in growth rate (in Magellanic penguins *Spheniscus magellanicus* [35]), together with a higher food conversion efficiency (in House sparrows *Paser domesticus* [36]) similar to in poultry. However, our understanding of the resilience of wild bird gut microbiomes to disruptions at the developmental stage is still limited. Such knowledge is important to understand the stability of host-microbe associations and consequences of microbiome disruptions in a current global environment where anthropogenic stresses continuously influence wild bird microbiomes [37–40].

As a step toward filling this gap, we explored the resilience of nestling gut microbiome to disruptions induced by antibiotics or probiotics during the brooding period in two sympatric altricial passerine (order Passeriformes) species: the Blue tit (BT: *Cyanistes caeruleus*) and the Great tit (GT: *Parus major*). First, we confirmed the influence of antibiotics and probiotics on gut microbiomes of adult birds (efficacy experiment). Then, we applied antibiotics and probiotics to nestlings in the wild and characterized the cloacal microbiomes throughout the brooding period with MiSeq amplicon sequencing of the bacterial 16s rRNA gene. We hypothesized that applying antibiotic or probiotic treatments would disrupt gut microbiomes, and that treated chicks would harbor less diverse and compositionally different microbiomes compared to control and non-treated chicks. However, if the microbiome transfer from the parents or the nest environment outweighs the disruptions caused by treatments, we expect the microbiomes of the chicks to differ less between treatments, prevailing the brood effect [14, 41].

## Materials and methods

### Study species

BTs and GTs are cavity-nesting passerines that readily accept nest-boxes for breeding [42]. This facilitates sample collection and, to some extent, homogenization of pre-treatment breeding parameters [43]. Both species are dimorphic and monogamous, with females building the nest, and laying and incubating the eggs, while both parents feed the brood [42, 44, 45]. BTs and GTs differ in body size [42] and therefore prey selection [46, 47], which is expected to affect gut microbiomes [23, 48].

### Testing the effects of antibiotics and probiotics (efficacy experiment)

The use of antibiotics alters the gut microbial communities of passerine birds compared to controls [36]. To confirm that our treatments influence gut microbiomes of tits, we conducted an efficacy experiment investigating the effect of commonly used broad-spectrum antibiotic Doxycycline (Doxygal 50mg/g) and probiotic *Lactobacillus fermentum* CCM7158 (Propigeon plv.) on ten GT adults, five per treatment. Experimental birds were mist-netted during winter 2020 (December-January) in Branišov forest (48°58’48”N, 14°25’23”E) in České Budějovice (Czech Republic). Birds were ringed, sexed, weighed (Table S1), and transported into a breeding room at the Faculty of Sciences, University of South Bohemia, České Budějovice (day 0). GTs were housed in individual cages, given fresh water (including vitamins (Acidomid exot®) three times per week) and fed a standard daily diet following a well-established protocol [18]. We surface-sterilized diet content with a UV lamp for 20 min before feeding it to the birds in order to minimize microbial input.

We handled the experimental GTs once per day in the early morning. Treatment started on day 1 and continued for three subsequent days (days 1-3). We supplied 0.5 mg of Doxygal or 6.7 mg of *Lactobacillus* probiotic per gram of body weight based on the weight at mist-netting, following product instructions, diluted in 0.25 ml of water for an adequate oral administration with a syringe. Birds were weighed (electronic scale 0.01 g) and cloacal swabs were taken (minitip Flocked Swab FLOQSwab® 501CS01) from day 1 (initial non-treated microbiome) to day 4, and later on day 8. Additionally, we collected two samples of diets to investigate the potential diet-associated microbial transfer. The handling of GTs and food was strictly done wearing nitrile gloves (cleaned with 70% ethanol between birds). After taking the last sample on day 8, we released GTs back to their original mist-netting location. We preserved swabs in 2 ml sterile vials filled with 100 μl of RNAlater® at -80°C until the DNA extractions.

### Manipulation of nestlings and sample collection (Field experiment)

The field experiment was conducted in a nest-box population in Branišov forest. All nest-boxes were inspected during the last week of April 2020. From that day on, BT and GT clutches at the incubation stage were checked daily until hatching. The first ten nest-boxes of each species that successfully hatched were assigned to the experiment (hatching date for each nestling = day 1) (Fig. 1). The first six hatchlings in each nest (average number of hatchlings per nest ± SD: BT = 11.3 ±1.25; GT = 8.9 ± 0.99) were randomly assigned in duplets to three treatments: antibiotic, probiotic, and control (day 1). The rest of the chicks in each nest-box remained untreated. We color-marked them with nontoxic pens for individual identification, took a cloacal swab and weighed them. In the field experiment we followed a slightly modified 16-day protocol compared with the efficacy experiment (Fig. 1). We changed the experimental procedure by administering the treatment every third day instead of daily, to avoid frequent disturbances in the nest. Control hatchlings were provided with water only (Table S2). Breeding parents were captured when entering their nest-boxes to feed the brood (day 10) by blocking the entrance of the nest-box. We sexed them based on plumage coloration and took a cloacal swab. Given the small size of the nestlings’ cloaca, all swabs were lubricated by immersing them in a vial filled with 2 ml of ultrapure water (IWA 20 IOL) just before use. We used separate vials per nest and visit and collected a water sample to control for potential microbiome transfer. Bird handling and storage of the cloacal samples was similar to the efficacy experiment.

**Fig. 1.**
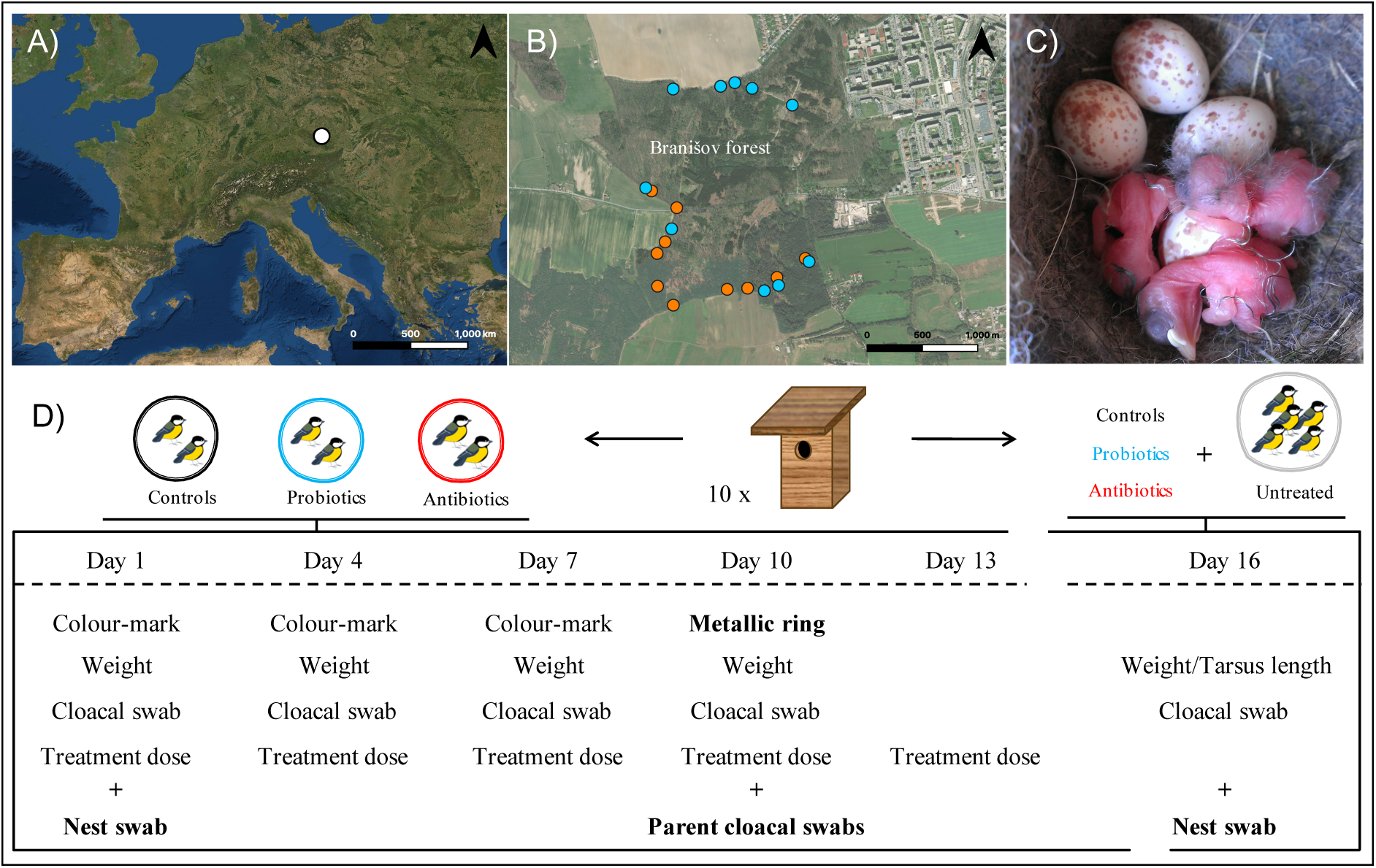
Location of the study site and schematic representation of the data collection in the field experiment. (A) The location of the study site (white dot), (B) the distribution of the experimental nest-boxes (blue dots refer to experimental Blue tit nests, orange dots to experimental Great tits nests), (C) first Great tit hatchlings in a nest, and (D) the experimental procedure on chicks in both Great and Blue tit nests (illustrations only represent Great tits). Day 1 = nestling hatching day.

### DNA extractions and MiSeq amplicon sequencing

DNA from cloacal swabs, food samples (from the efficacy experiment) and water samples used to lubricate swabs (field experiment) were extracted using Qiagen DNeasy blood and tissue kit® (Hilden, Germany) following an already validated protocol [49]. The presence of bacterial DNA in samples was validated using primers (SA511 and SB701) targeting bacterial 16S rRNA gene and samples were sequenced on an Illumina MiSeq platform at the Microbiome Core at University of Michigan.

### Data analyses

MiSeq amplicon sequences were cleaned and aligned using the DADA2 pipeline [50] within QIIME2 [51]. Sequences were clustered into amplicon sequence variants (ASVs) at 100% similarity and assigned to taxonomy using the SILVA 132 bacterial database [52]. All chimeric, archaeal, mitochondrial and chloroplast sequences were removed following the QIIME pipeline. We detected contamination in the water samples used to lubricate the cloacal swabs. These sequences were consistently found across samples and were removed from the full dataset. A rooted bacterial phylogeny was acquired using the *align-to-tree-mafft-fasttree* command in QIIME2. We also removed samples with less than 3,000 sequences from further analyses. Subsequent analyses were conducted separately for the efficacy and the field experiment.

Each dataset was rarefied using the sample with the smallest number of sequences (efficacy experiment: 5,158 and field experiment: 3,037) to correct for differences in sequencing depths using *rarefy_even_depth* function in the phyloseq package [53] (Tables S3 and S4) and subsequent analyses were conducted in R 4.0 [54]. Using the *diversity* function in the microbiome package [55], we calculated multiple alpha diversity matrices: observed ASV richness, Shannon’s diversity index, and relative dominance (relative abundance of the most abundant bacterial taxa). We further calculated Faith’s phylogenetic diversity of microbial communities using the picante package [56].

The effect of experimental treatments on body mass and alpha diversities in the efficacy experiment was assessed by building linear mixed-effect models (LMMs) using the lme4 package [57]. We used treatment (antibiotics or probiotics), day of experiment (quadratic term via *poly* function), sex (male or female), and the interaction between treatment and day of experiment as fixed explanatory variables. Bird identity was used as a random intercept variable.

In the field experiment, we conducted separate analyses for BT and GT chicks. We investigated body mass and alpha diversities from day 1 to 16, by building models containing treatment (control, antibiotics or probiotics) and the day of the experiment (quadratic term via *poly* function) as fixed explanatory variables. We added brood identity as a variable with random intercept because of its grouping nature and modelled the repeatability of chick microbiome sampling with random slopes within day of experiment. We assessed tarsus length, as a proxy for size, at day 16, between all chick groups by building a LMM using the experimental category as a fixed factor (untreated, control, antibiotics, probiotics), and brood identity as a random variable. We additionally compared alpha diversity indexes between all chick groups, adults and the nest microbial environment at day 16 (last day of experiment) using group as a fixed factor (untreated, control, antibiotics, probiotics, male, female and nest) and brood identity as a random variable. We also conducted analyses comparing alpha diversity indexes between chicks (pulling together the three experimental treatments) and nests at day 1 and at day 16 using group (nest and chick) and the day of the experiment as fixed factors, including their interaction, and brood identity as a random variable. Continuous explanatory variables were centered and, when necessary, the data were log or square root transformed (see Tables S5 and S6). We used the *r.squaredGLMM* function from MuMIn package [58] to compute R^2^ values [59].

To investigate bacterial community structures (beta diversities) we used Bray-Curtis and weighted UniFrac (accounting for bacterial phylogeny) distances and visualized using non-matric multidimensional scaling (NMDS) and principal coordinate analysis (PCoA) plots. The influence of different treatment types and sample types were assessed using permutational multivariate analyses of variance (PERMANOVAs) with the *adonis2* function in the vegan package [60] with the “by” parameter set to “margin” to assess the marginal effect of the tested variables. Pairwise differences in microbial communities were investigated using the pairwiseAdonis wrapper package [61]. To investigate the effect of treatments on associations between microbes in the efficacy experiment, we calculated microbial co-occurrence networks with the *trans_network* function in microeco package [62] using the SparCC method from the SpiecEasi package [63]. We filtered out ASVs with abundances below 1% from the data set and used 100 SparCC simulations. The network properties were calculated with the igraph package [64] and visualized using Gephi [65].

Similarly to alpha diversities, field experiment beta diversities were analyzed separately by host species, for manipulated chicks and for final day samples. To tease apart the parental transmission of microbiomes to chicks, we analyzed microbiomes between adults and 16-day old chicks (treated and untreated). We first evaluated the parental and environmental transfer of ASVs through characterizing shared and unique ASVs between adults (males and females), chicks, and nest environment using *UpSet* plots in the UpSetR package [66]. Secondly, we investigated the influence of parental and nest microbiomes on core microbiomes (consistent bacterial taxa) of nestlings using the *core* function in the microbiome package [55]. We assigned an ASV to the core if the ASV was found in abundances of a minimum of 0.001% across >50% of the samples in the same treatment group. We also examined the transfer of microbiomes from the nest through comparing the nest microbiomes with chicks on day 1 (the day a chick hatched), under the assumption that recently hatched chicks do not strongly influence nest microbiomes, but reversely, the nest environment may influence the bacterial communities in the chicks. We conducted similar analyses between chick and nest microbiomes of day 16 to investigate whether chick microbiomes converge similar to nest microbiomes at the end of the brooding period. We further investigated the influence of the distance between nests on microbiome similarity of chicks using a *mantel test* in the vegan package [60], to evaluate whether the proximity of nests (i.e., similar environmental conditions and diet availability) influence chick microbiomes. We used nest-box GPS coordinates (Table S7) to calculate the distances between them, using the *st_distance* function from the sf package [67]. For this analysis we averaged the chick microbiomes from the final day from each nest.

## Results

### Notable disruptions to adult gut microbiomes in the efficacy experiment

In the efficacy experiment, 47 of the 50 samples passed the quality filtering steps and these samples contained overall 1 799 504 (average ± SD = 38 287 ± 25 372) bacterial sequences. These sequences were assigned to 4 886 ASVs. GTs lost body mass after 8 days in captivity (average ± SD = 12.6 g ± 4.97%, n = 10), which is expected from the adaptation to laboratory conditions [68, 69]. Mass loss was only associated with the number of days since the beginning of the experiment (estimate ± SE = -2.71 ± 0.282, t = 9.623, *p* <0.001), independent of treatment (estimate ± SE = -0.05 ± 0.270, t = -0.196, *p* = 0.850) (Fig. 2A). Observed ASV richness and Faith’s phylogenetic diversity declined similarly in both antibiotic and probiotic treated individuals during the experiment (Figs. 2B, S1, and Table S3) while we did not detect any temporal pattern in Shannon’s diversity index or relative dominance (Fig. S1, Table S3).

**Fig. 2.**
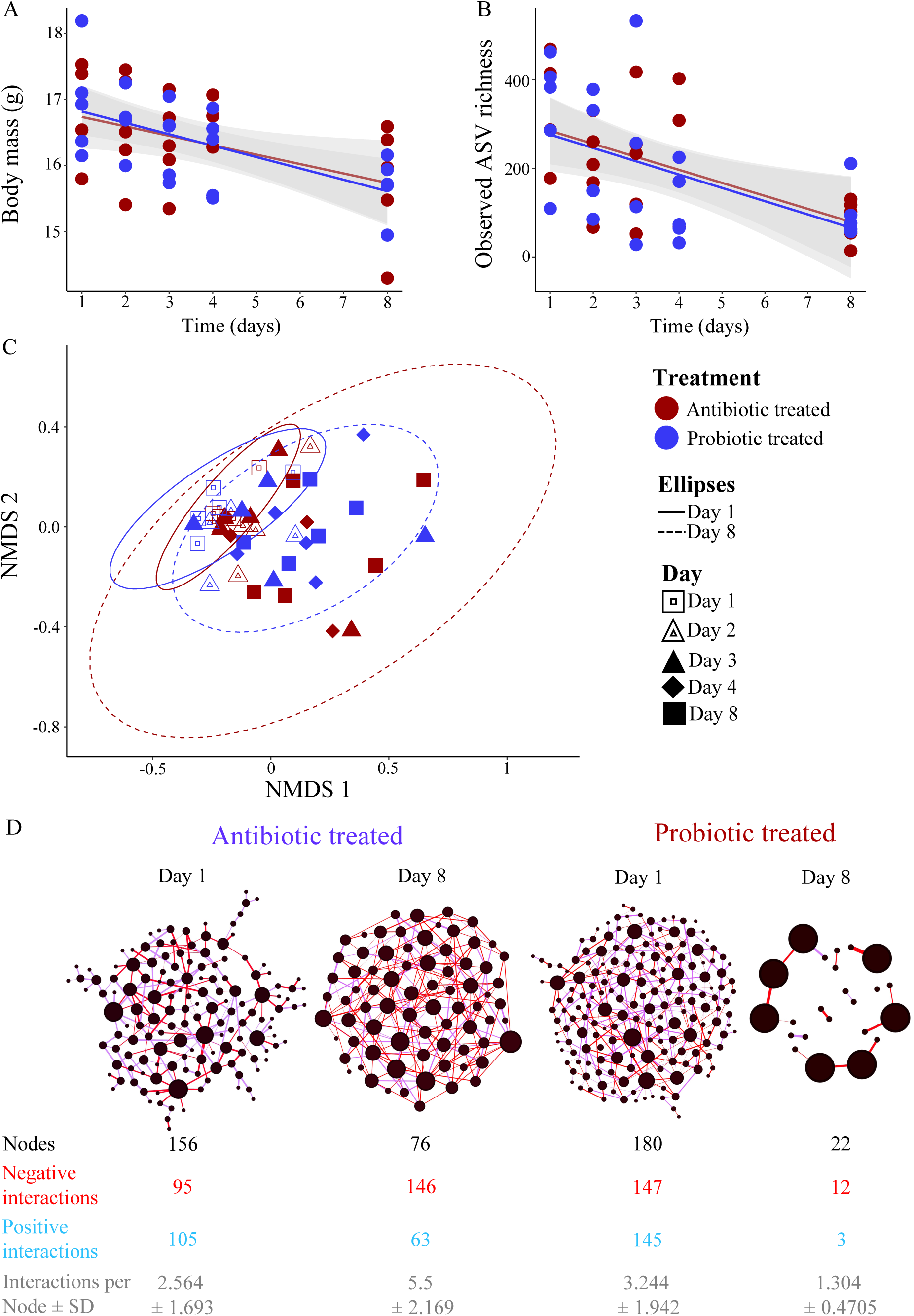
Antibiotic and probiotic treatments influence host body mass, and alpha and beta diversities of microbiomes during the efficacy experiment. (A) Decline in body mass during the efficacy experiment in both antibiotic and probiotic treated individuals. Standard errors for regression lines are indicated with grey shaded area. (B) Observed ASV richness of gut microbial communities decreased at the end of the treatment period compared to initial microbiomes. (C) Non-metric multidimensional scaling plot (NMDS) of the gut microbial communities of antibiotic and probiotic treated individuals (stress = 0.243). Individual microbiome variation was lower at the beginning of the experiment (Day 1: solid-line ellipse) in both treatment groups compared to the microbial variation at the end of the treatment (Day 8: dashed-line ellipse). Ellipses represent 95% confident intervals of the data. (D) Microbial co-occurrence networks of initial microbiomes (Day 1) and at the last sampling day (Day 8) after the treatments. Nodes represent individual ASVs, and size correspond to the degree of the ASV (number of interactions each ASV have with other ASVs) in each network. Edges represent whether association are positive (blue) or negative (red). Network attributes are given below each network.

At the phylum level, Proteobacteria dominated the microbiomes (50.2%) followed by Bacteroidetes (17.2%), Tenericutes (27.5%), Actinobacteria (9.6%) and Firmicutes (5.2%) (Fig. S2). In both antibiotic and probiotic treated individuals, the relative abundance of Proteobacteria (Antibiotic: Initial = 56.2%, last day = 38.7%; Probiotic: Initial = 53.2%, last day = 25.3%) and Bacteroidetes (Antibiotic: Initial = 23.5%, last day = 12.5%; Probiotic: Initial = 22.8%, last day = 2.9%) decreased during the treatment time, while Tenericutes increased notably (Antibiotic: Initial = 4.3%, last day = 28.4%; Probiotic: Initial = 1.3%, last day = 60.7%) (Fig. S2). In the probiotic treatment, the relative abundance of Firmicutes had increased from the first to the last day, potentially influenced by the inoculation of lactic-acid bacteria (Fig. S2). Microbial community compositions measured with either distance matrices were not significantly different between sampling days for both treatment groups (antibiotic _(Bray-Curtis)_: F = 1.054, R^2^ = 0.1898, *p* = 0.2516, antibiotic _(UniFrac)_: F = 1.227, R^2^ = 0.2143, *p* = 0.1139, probiotic _(Bray-Curtis)_: F = 1.196, R^2^ = 0.2012, *p* = 0.1316, probiotic _(UniFrac)_: F = 1.036, R^2^ = 0.1791, *p* = 0.3719; Fig. 2C). However, microbiomes on day 1 exhibited reduced interspecific variation compared to the last day in both treatment groups (Fig 2C), indicating that antibiotic and probiotic treatments increase variability in microbiomes between individuals.

Microbial network analyses confirmed the effects of antibiotic and probiotic treatment on changing co-occurrence patterns of ASVs between days 1 and 8 (Fig. 1D). The number of ASVs in networks reduced during the period, while the proportion of negative associations between ASVs increased in both groups (Fig. 1D). Overall, this indicates that the influence of antibiotic and probiotic treatments on alpha and beta diversities of microbiomes lead to notable changes in structure of microbial networks.

We observed the presence of a few food-borne ASVs in the gut communities (Fig. S3A and B), but these ASVs were abundant, as community composition of food microbiomes were similar to some of the gut microbiomes (Fig. S3C). Removal of these food-borne microbes from the dataset did not influence overall community compositions or effects of antibiotic or probiotic treatment, so we retained them in the dataset.

### Treatment does not affect gut microbiomes of manipulated chicks in the wild

In the field manipulation experiment, from chicks, we acquired 4 356 709 bacterial sequences from BTs (n = 203, average ± SD: 21 461 ± 10 158) and 6 179 421 sequences from GTs (n = 257, average ± SD: 24 044 ± 11 794), and these sequences were assigned to 14 309 ASVs in BT and 18 796 ASVs in GT samples. Alpha diversity indexes of microbiomes increased overall during chick development in both species (except for the relative dominance index) with major changes occurring between day 1 and day 7 (Figs. 3, S4 and Table S4). Alpha diversity indexes showed a clear negative quadratic effect, stabilizing between day 10 and 16, following the decrease in body growth rate (Figs. 3 and S4). We did not find a clear effect of treatment on the diversity matrices in BTs, but GTs showed a trend for higher observed ASV richness and Faith’s phylogenetic diversity in antibiotic treated chicks than in controls (Fig. 3, Table S4, and S5).

**Fig. 3.**
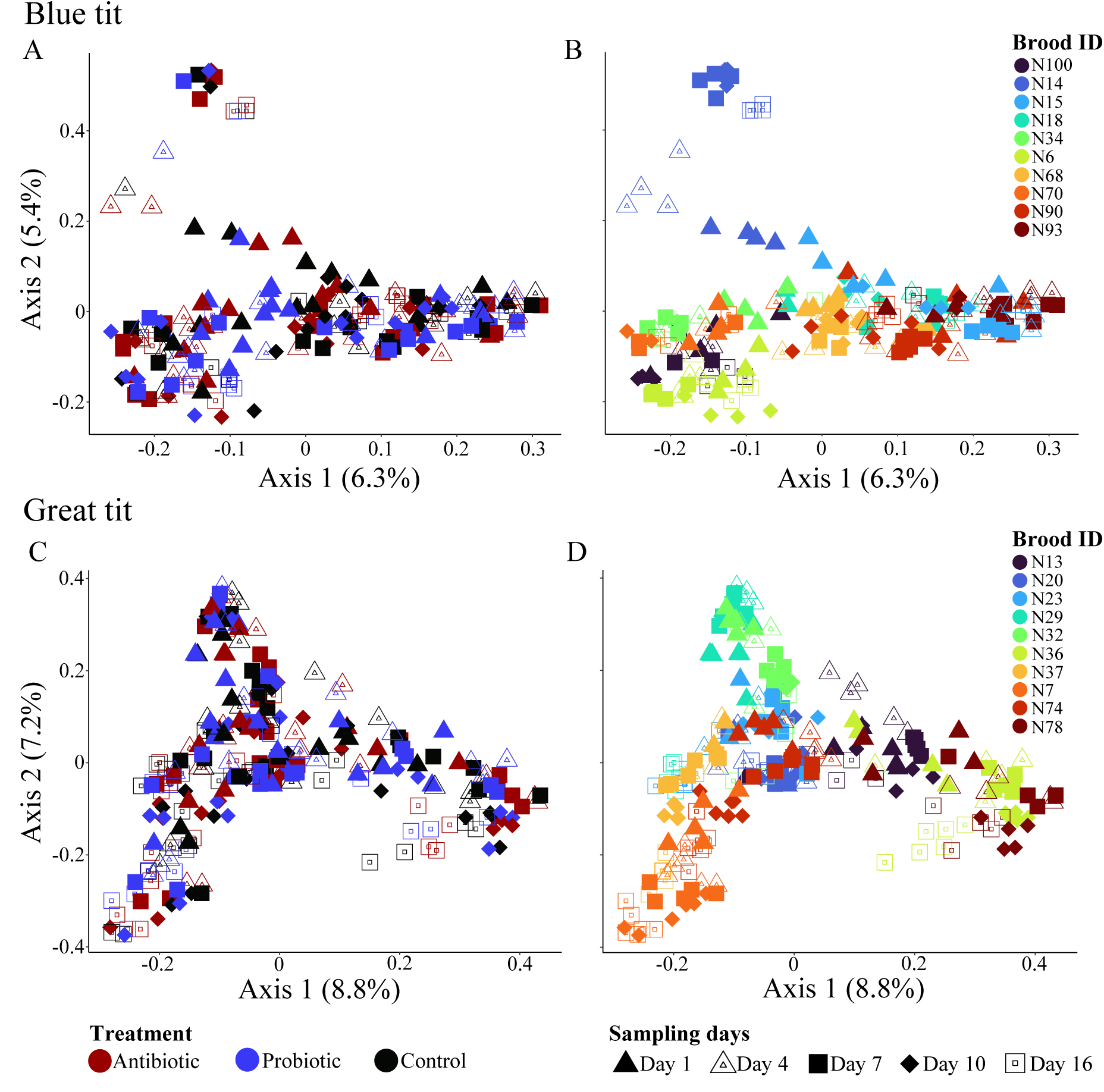
Growth rate and gut microbial alpha diversities did not differ between chicks in different treatment groups. Body mass (A), observed ASV richness (B), and Faith’s phylogenetic diversity (C) of microbiomes of manipulated chicks during the sampling period. Gray areas around the trend lines represent standard errors. Data points are colored according to the treatment.

The increase in body mass during development was similar in antibiotic and probiotic treated chicks as well as controls for both species (Fig. 3, Table S6). However, on day 16, GT probiotic treated chicks tended to be larger than untreated (estimate ± SE = 0.27 ± 0.140, t-value = 1.933, *p* = 0.057) and antibiotic treated chicks (Table S7).

The microbiomes of manipulated chicks were dominated by Proteobacteria (BT: 37.6%, GT: 38.1%), Firmicutes (BT: 19.1%, GT: 19.3%), Actinobacteria (BT: 18.1%, GT: 18.0%) and Bacteroidetes (BT: 17.8%, GT: 15.8%) bacterial phyla and the relative abundance of these phyla did not differ between days nor treatments (Fig. S5). The composition of microbiomes (beta diversity) was strongly affected by brood identity and sampling day (Fig. 4, Table 1), irrespective of the distance matrix used. Antibiotic or probiotic treatments did not strongly influence microbiome compositions in developing chicks (Table 1).

**Fig. 4.**
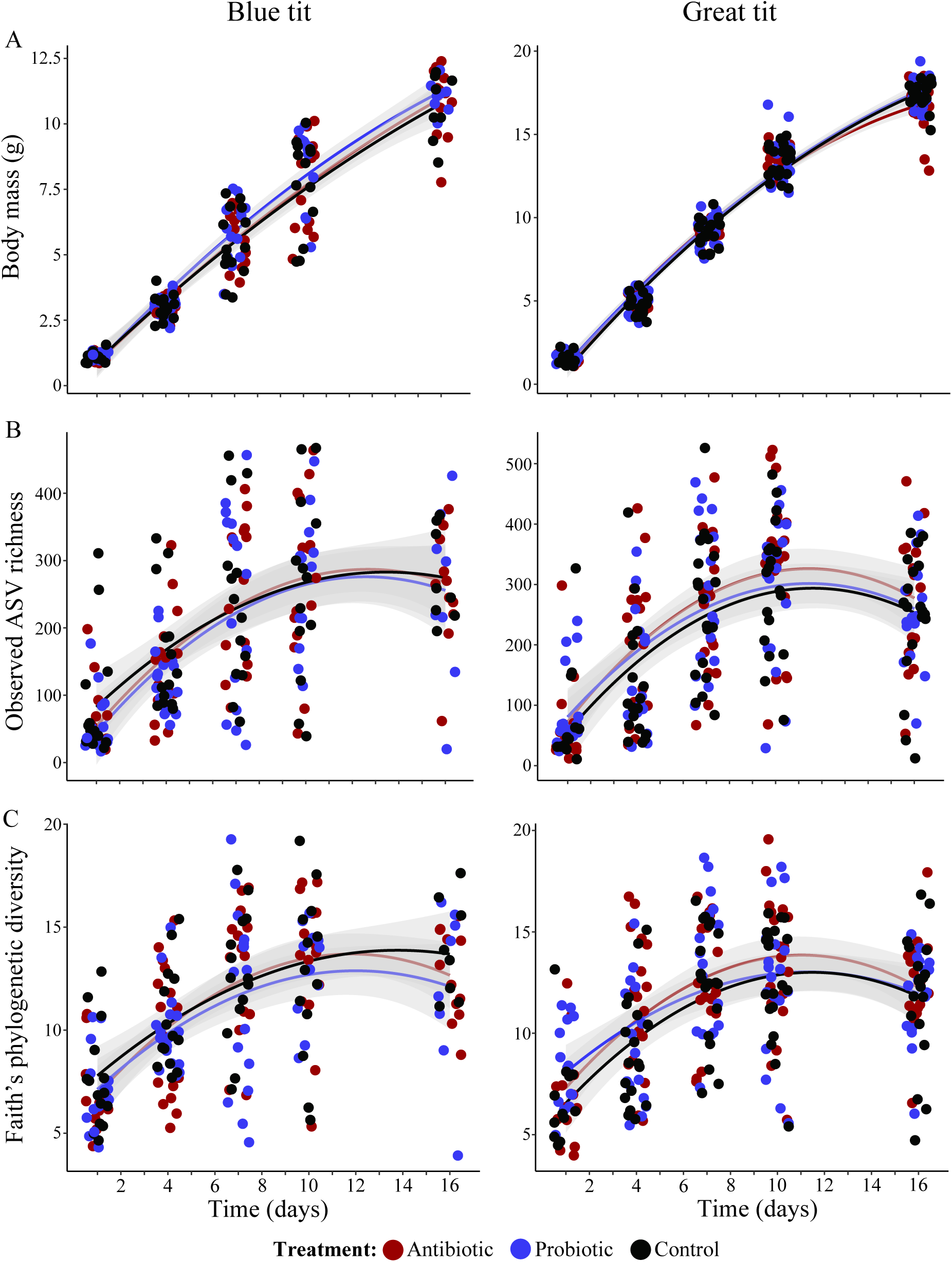
Nest environment has a stronger effect than antibiotic/probiotic treatments on shaping the gut microbiomes of developing chicks. Microbial communities of manipulated chicks of Great (A, B) and Blue (C, D) tits. Individuals in A and C are colored according to the treatment, while individuals in B and D are colored according to nest. Shapes indicate day of sampling across all four plots.

**Table 1.**
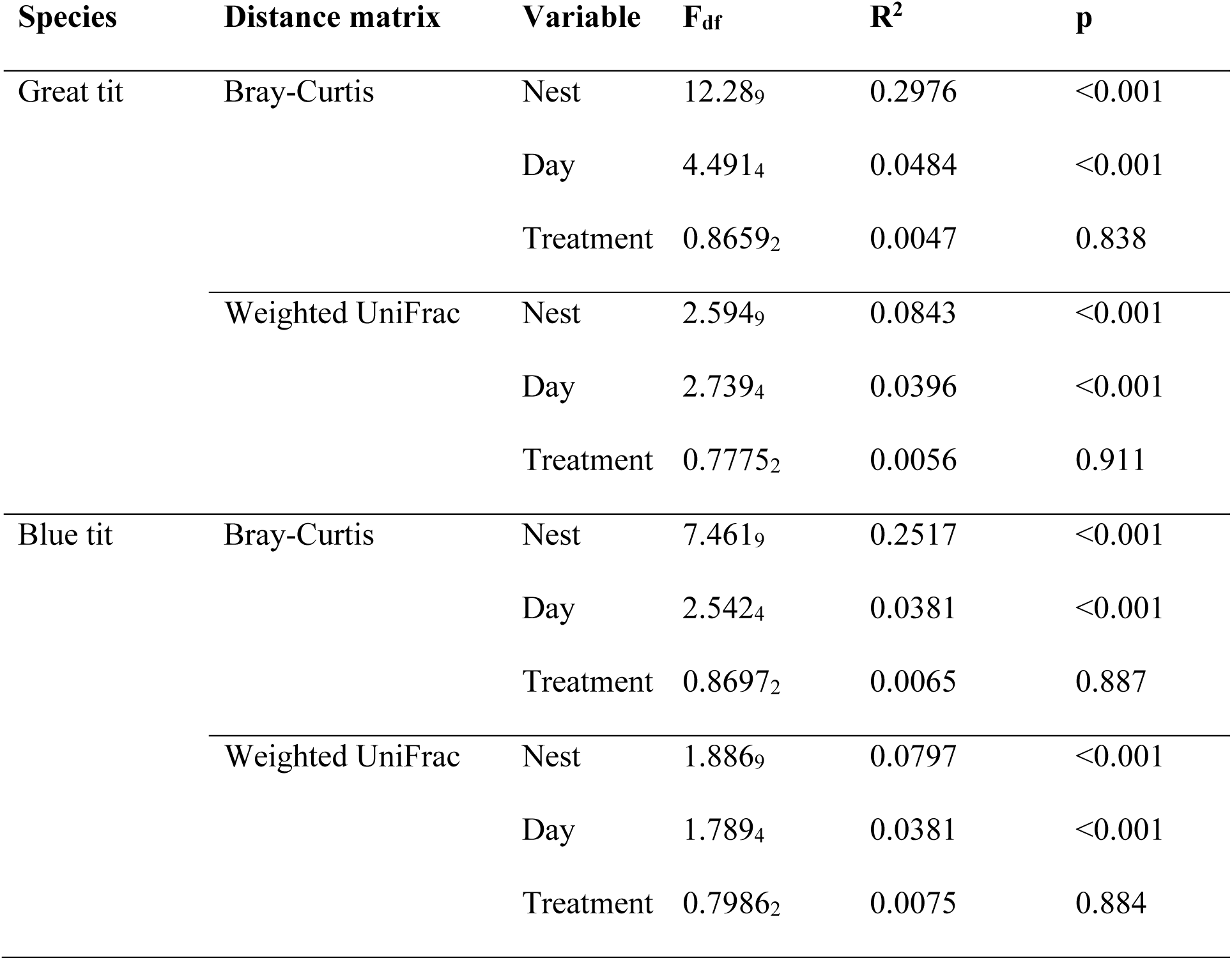
Influence of nest, treatment day and treatment on the composition of gut microbial communities in the manipulated chicks based on permutational multivariate analyses of variance (PERMNOVAs) tests. Analyses were conducted through measuring community composition using both Bray-Curtis and weighted UniFrac distances.

| Species | Distance matrix | Variable | F<sub>df</sub> | R<sup>2</sup> | p |
| --- | --- | --- | --- | --- | --- |
| Great tit | Bray-Curtis | Nest | 12.28<sub>9</sub> | 0.2976 | <0.001 |
|  |  | Day | 4.491<sub>4</sub> | 0.0484 | <0.001 |
|  |  | Treatment | 0.8659<sub>2</sub> | 0.0047 | 0.838 |
|  | Weighted UniFrac | Nest | 2.594<sub>9</sub> | 0.0843 | <0.001 |
|  |  | Day | 2.739<sub>4</sub> | 0.0396 | <0.001 |
|  |  | Treatment | 0.7775<sub>2</sub> | 0.0056 | 0.911 |
| Blue tit | Bray-Curtis | Nest | 7.461<sub>9</sub> | 0.2517 | <0.001 |
|  |  | Day | 2.542<sub>4</sub> | 0.0381 | <0.001 |
|  |  | Treatment | 0.8697<sub>2</sub> | 0.0065 | 0.887 |
|  | Weighted UniFrac | Nest | 1.886<sub>9</sub> | 0.0797 | <0.001 |
|  |  | Day | 1.789<sub>4</sub> | 0.0381 | <0.001 |
|  |  | Treatment | 0.7986<sub>2</sub> | 0.0075 | 0.884 |

### Maternal microbial transfer is important for structuring chick microbiomes

From adults, we acquired 283 044 sequences in BTs (males (n = 6): 21 265 ± 7 832; females (n = 8): 19 432 ± 10 328) and 272 345 sequences in GTs (males (n = 10): 19 342 ± 8 496; females (n = 4): 19 729 ± 8 496). Overall, the phylogenetic diversity of microbiomes did not differ between chicks at day 16 and adults in both species (Fig. 5A, and S6, Table S8). GT adult male microbiomes showed a lower bacterial richness and Shannon’s diversity index, compared to chicks (Fig. 5A, and S6, Table S8). For adult BTs, we only found a lower Shannon’s diversity index of both males and females than chicks (Fig. 5A, and S6, Table S8).

**Fig. 5.**
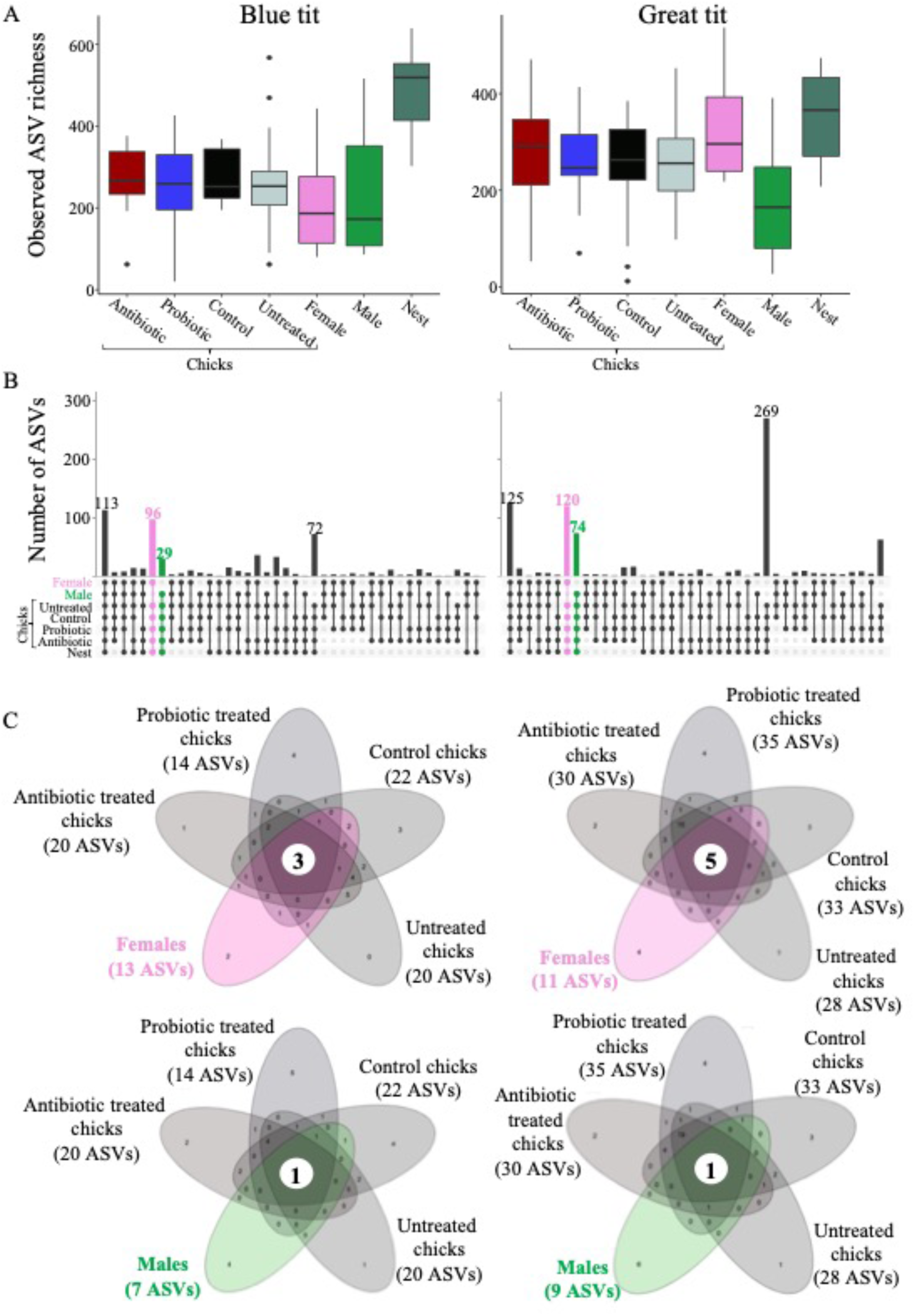
Alpha diversities of parent microbiomes did not differ from chicks, but maternal microbiomes contributed notably to the composition of chick microbiomes. (A) Box plots depicting the observed ASV richness in chicks and adults. (B) Upset plots showing the number of shared and unique ASVs found between chicks and adults. Unique ASVs found between chicks and female or male are indicated with colored bars. (C) Flower plots depicting the shared core microbiome between chicks and adults.

Of the bacterial ASVs shared between adults and chicks, females shared a higher number of unique ASVs with chicks (BT: 96, GT: 120) than males (BT: 29, GT: 74), indicating a strong effect of maternal microbiome transfer to chicks (Fig. 5B). Overall, core microbiomes were small in all groups, with chicks harboring a larger core microbiome than adults (Fig. 5C). The smaller core microbiomes in adults could be driven by environmental and dietary impacts on microbial variation and/or that fewer adults were examined than chicks [18, 70]. Despite small core microbiomes, chicks shared more core taxa with females (BT: 3 ASVs and GT: 5 ASVs) than males (1 ASV in both BT and GT) (Fig. 5C), underscoring the stronger maternal than paternal effect.

Adult microbiome compositions differed from the microbiomes of chicks on the last sampling day (Fig. S7A). Relative abundance of major bacterial phyla, such as Proteobacteria, Actinobacteria and Bacteroidetes, were comparable between adults and chicks (Fig. S7A). However, in both bird species, adult birds harbored a larger relative proportion of Tenericutes (BT females: 31.8%; males: 35.8%, chicks: 0.4%; GT females: 26.8%; males: 26.6%, chicks: 1.2%), and a lower relative proportion of Firmicutes (BT females: 8.9%; males: 2.5%, chicks: 11.4%; GT females: 6.6%; males: 3.5%, chicks: 25.4%) compared to all chicks on the last day (Fig. S7A).

Microbiome composition was significant different between treatment groups (PERMANOVA_10,000 permutations_: BT: F_6_ = 1.408, R^2^ = 0.1166, *p* < 0.0001 and GT: F_6_ = 1.728, R^2^ = 0.0975, *p* < 0.0001) (Fig. S7B and C). The pair-wise comparisons confirmed the reduced influence of paternal microbiomes on the composition of chick microbiomes, as we observed differences in microbial community composition between males and chicks (Table S9). Female microbiome composition did not differ from manipulated chicks but did differ from untreated chicks (Table S9). This suggests that disruptions to the microbiomes increased transfer of maternal microbes to treated chicks. Taken together, these results indicate that maternal transfer of microbes to developing chicks counter disturbances to developing microbiomes, but that only a subset of maternal microbes establish in chick guts.

### Environmental transfer of microbiomes

From the nest environment, we acquired 337 615 (n = 10; average ± SD = 33 762 ± 15 502) bacterial sequences from BTs and 323 275 (n = 10; 32 328 ± 13 041) from GTs on day 1, and 200 844 sequences from BTs (n = 8; 25 106 ± 4 372) and 252 088 (n = 10; 25 209 ± 6 478) from GTs on the day 16. Bacterial richness, Shannon’s diversity and phylogenetic diversity of nest microbiomes did not differ between the sampling times for GTs, but BT nests increased in bacterial richness and phylogenetic diversity over time (Figs. 6, S8A and B, and Table S10). Nest beta diversity did not differ between days 1 and 16 for neither bird species (Fig. S8C).

**Fig. 6.**
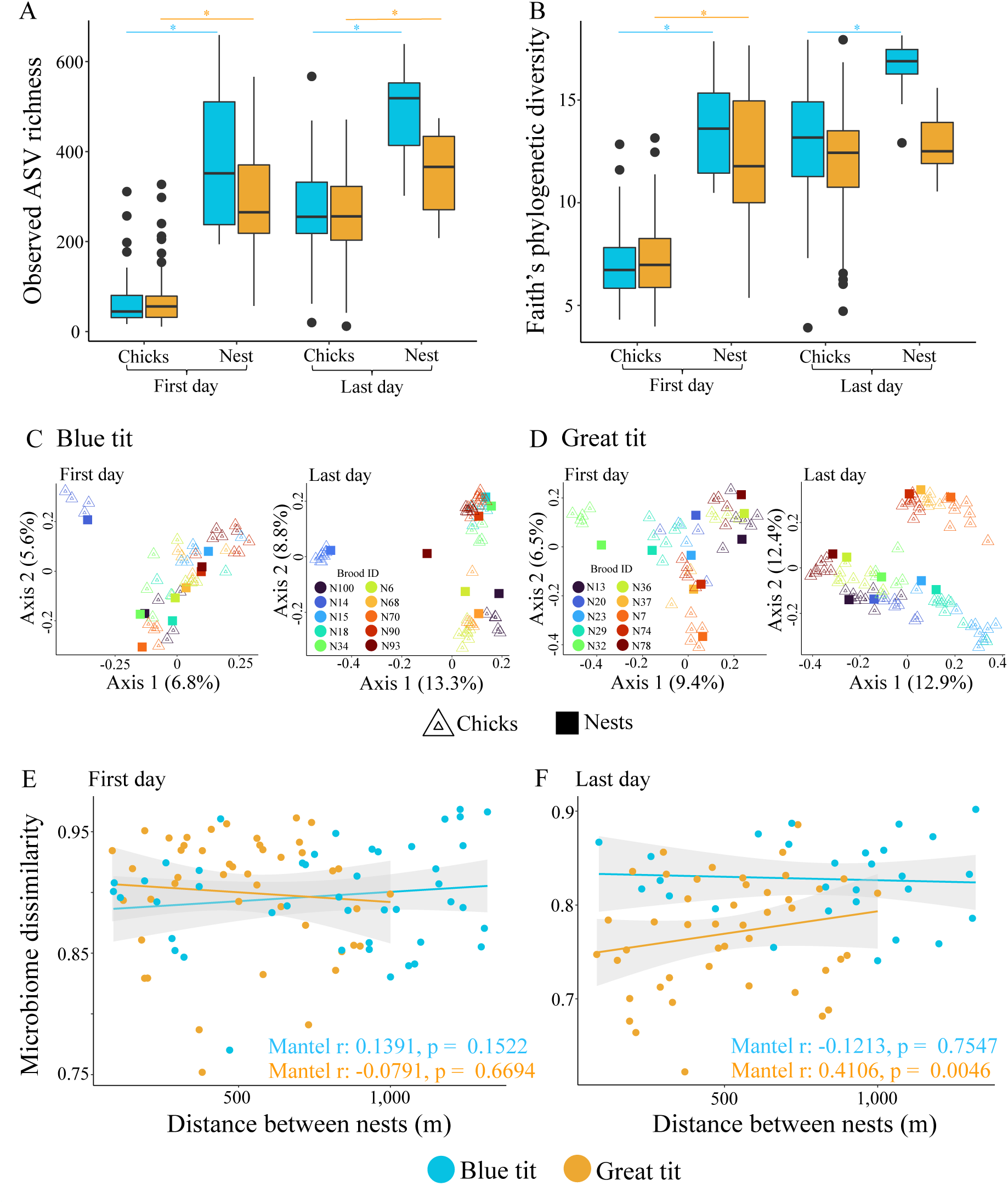
Nest microbiomes tend to influence cloacal microbiomes of chicks. Observed ASV richness (A) and Faith’s phylogenetic diversity (B) of chick and nest microbiomes during first and last sampling days. Statistical differences are indicated with asterisks. Ordination plot depicting the compositional differences (measured with Bray-Curtis distances) in chick and nest microbiomes during first and last sampling days of Blue (C) and Great (D) tits. Association between distance among nests and average chick microbiome per nest in first (E) and last (F) sampling days. Mantel test statistics are given within each graph.

Alpha diversities were significantly higher in nest microbiomes than chick gut microbiomes on day 1 (Fig. 6A and B and Table S10) and on day 16, except for phylogenetic diversity in GTs (Fig. 6A and B and Table S10). We observed increased microbial diversity in chicks towards the end of the brooding period compared to the hatching day. Microbial diversity in chicks on day 16 (just before fledging) became similar to their nests. The microbiome composition of BTs was significantly different between nests and one day old chicks (PERMANOVA_10,000 permutations_: F_1_ = 1.581, R^2^ = 0.0307, *p* = 0.0013) (Fig. 6C). This was not the case for GTs (PERMANOVA_10,000 permutations_: F_1_ = 1.078, R^2^ = 0.0211, *p* = 0.2812) (Fig. 6D). Despite the significant difference between community composition of day 1 chicks and nest microbiomes in BTs, the visual inspection of ordination plots indicated that this is driven by higher variability in chick microbiomes than nest microbiomes (Fig. 6C).

The similarity in average chick microbiome composition on day 1 (per brood) was not associated with distance between nests (Fig. 6E). This was retained in BTs until the last day of sampling (day 16) but there was a strong positive correlation between nest distance and gut microbiome dissimilarity in GTs on day 16 (Fig. 6F), suggesting that the effect of environment varies between species.

## Discussion

Here we investigated the influence of continuous disruption on the establishment of the gut microbiomes during development in two altricial wild bird species. Despite the influence of antibiotic and probiotic treatments on gut microbiomes of adult birds, treatments did not negatively impact the diversity and composition of the gut microbiomes of nestlings during development. Consistent with previous studies [14, 26–28, 41], we observed a strong brood effect, driven by the continuous transfer of microbes from the parents and the potential transfer of microbes from the environment (e.g., nest and diet associated microbes). This underlines the importance of environmental and parental transfers, including rescue after disruption, for microbiomes in chicks with plausible influence on both their health and fitness later in life.

The strong within-brood similarity in chick gut microbiomes underlines the importance of parents in shaping offspring gut communities during the brooding period, through direct (e.g., feeding events [71, 72]) and indirect (e.g., accumulation of parental microbes in the nest itself [14, 15]) microbial transfer. Parents usually take turns feeding the brood (10 to 40 individual feeding events per hour [46, 71–73]), with similar frequencies in males and females [74, 75].

This should ensure a continuous inoculation of parental microbes to chicks, leading to strong convergence of offspring gut microbiome within a brood, while counteracting disruptions to gut microbiomes during development. However, despite bi-parental care, we observed a stronger effect of maternal than paternal microbiomes on chick gut microbiomes (see also [27]). Females spend more time than males during nest building [42, 76], egg laying [77–79], incubation [80–82], and brooding [42, 83, 84], which could lead to a maternally biased shedding of microbes that can be acquired by the chicks. Indirect transfer of maternal microbes via nest environment has also been shown in Zebra finch chicks (*Taeniopygia guttata*) [15]. Similar mechanisms are likely in both tit species we studied, as we observed comparably stable microbiomes within nests during the brooding period (Figs. 6A, B and S8), higher similarity of maternal and nest microbiomes than paternal and nest microbiomes (Fig. 5B and Table S8), and higher levels of microbiome sharing and convergence between chicks and nests during the brooding period (Fig. 5B).

The direct and indirect transfer of maternal microbiomes is likely essential for naturally developing chick microbiomes, as they may lose some gut symbionts due to diet and habitat changes, and during infections with natural pathogens or ones associated with anthropogenic activities [16, 23, 38, 85]. Skewed maternal microbial transfer may reduce competition between parental microbial symbionts sharing the same niches within offspring guts, with potential deleterious effects to chicks [86, 87]. However, male microbial symbionts are not completely lost during generational transmission, indicating that colonization of males does not necessarily mean a dead end for microbes. This may thus reflect a bet-hedging situation for optimal access to important symbionts, secured through biparental transfer. However, it appears more likely that the maternal-biased microbial transfer is derived mainly from the dominant role of females during the breeding period, implying that species with more equal biparental contribution throughout the nest building, incubation and brooding periods should also exhibit more equal transfer of parental microbes to the next generation.

In GTs, nests located further from each other were more different in gut microbiome structure at the end of the brooding period, highlighting the joint impact of the microhabitat and diet on gut microbiomes. Surprisingly, we did not detect such an association in BTs. Previous work has shown that differences in habitat composition influence wild bird gut microbiomes [23], specifically, the microbiome similarity between prey and predator (caterpillar – tit) is higher when the prey is captured closer to the nest-box [17]. Our observed interspecific differences could originate from differential foraging behaviors or habitat quality of the proximal environment. Tits usually forage within a 25 m radius of the nest-box and increase travelling distances when resources are scarce [88]. GTs are larger and dominant over BTs in competing for nest-boxes [89, 90], which may lead GTs to select nest-boxes in higher-quality habitat patches compared to BTs. Consequently, GTs could forage closer to the nest, which reinforces the association between gut microbiomes and nest-box location. As a result, BT nests might be located in lower-quality habitat patches associated with longer foraging distances [73, 91], reducing the strength of the gut microbiome–location association. Alternatively, or in conjunction with habitat quality, GT is a more generalist species than BT [47, 92], and may be able to exploit multiple food resources near the nest. The more specialist BTs would have to forage further away if their preferred prey is scarce nearby, leading to a reduced influence of nest location on chick gut microbiomes. Overall, this indicates that nest location can also influence the gut microbiome composition of developing chicks, but this effect may depend on prey preference and foraging behavior of the species.

## Conclusions

Disruptions to early-life establishment of gut microbiomes can have negative consequences for the development and fitness of animal hosts. Our gut microbiome manipulation study in natural environments highlights the resilience, yielded by parental feeding and environmental acquisition of microbes from nests, to disruptions of gut microbiomes during early life in two altricial bird species. Despite the counteracting effects of continuous transfer of maternal gut microbiomes during chick development, the influence of nest and diet-associated microbes indicate that chick microbiomes are still vulnerable to the introduction of new bacterial symbionts during brooding. If pathogenic, these newly arriving symbionts occupying niches opened-up by gut disruptions may hinder natural host microbial associations, affecting host development and compromising health. Altogether, our findings indicate that maternal-driven transfer of microbial symbionts is important for the establishment and stability of chick microbiomes, potentially affecting long-term associations between avian hosts and their gut symbionts.

## Ethics declaration

All the necessary permits were obtained for this experimental project: licence no. 1004 issued by the National Museum in Prague to capture wild birds, licence no. 43873/2019-MZE-18134 issued by the Czech Ministry of Agriculture to house wild birds and licence no. MZP/2020/630/1544 granted by the Czech Ministry of Environment to conduct behavioural experiments with wild birds.

## Availability of data and material

Microbiome sequences are submitted to Sequence Read Archive database in GenBank (Efficacy experiment: PRJNA800248, Field experiment: PRJNA800611), and accession numbers of samples are available in Zenodo (doi: 10.5281/zenodo.6174091).

## Acknowledgements

This project and K.S., D.D-M., I.F. and I.K. were financially supported by the European Research Council Starting Grant BABE 805189. KAJ is grateful for the financial support received from the Villum Foundation (Young Investigator Programme, project no. 15560) and the Carlsberg Foundation (Distinguished Associate Professor Fellowship no. CF17-0248). We thank Dr. Petr Veselý and Dr. Michaela Syrová for their help mist-netting Great tits, and Dr. Pável Matos-Maraví for his help with housing birds.

## Author contributions

The study was conceived by K.S., K.B. and D.D-M., the efficacy and field experiment were carried out by I.F. and D.D-M.; I.K. carried out the laboratory work and K.B. and D.D-M. analyzed the data. K.B and D.D-M. led the writing of the manuscript, with inputs from I.F and I.K and critical contributions from K.A.J., M.P, and K.S.

## References

1. Heijtz RD, Wang S, Anuar F, Qian Y, Bjorkholm B, Samuelsson A, et al. Normal gut microbiota modulates brain development and behavior. Proc Natl Acad Sci USA 2011; 108: 3047–3052.

2. Alberdi A, Aizpurua O, Bohmann K, Zepeda-Mendoza ML, Gilbert MTP. Do vertebrate gut metagenomes confer rapid ecological adaptation? Trends Ecol Evol 2016; 31: 689–699.

3. Macke E, Tasiemski A, Massol F, Callens M, Decaestecker E. Life history and eco-evolutionary dynamics in light of the gut microbiota. Oikos 2017; 126: 508–531.

4. Davidson GL, Raulo A, Knowles SCL. Identifying microbiome-mediated behaviour in wild vertebrates. Trends Ecol Evol 2020; 35: 972–980.

5. Bodawatta KH, Hird SM, Grond K, Poulsen M, Jønsson KA. Avian gut microbiomes taking flight. Trends Microbiol 2022; 30: 268–280.

6. Jacob S, Parthuisot N, Vallat A, Ramon-Portugal F, Helfenstein F, Heeb P. Microbiome affects egg carotenoid investment, nestling development and adult oxidative costs of reproduction in Great tits. Funct Ecol 2015; 29: 1048–1058.

7. Davidson GL, Wiley N, Cooke AC, Johnson CN, Fouhy F, Reichert MS, et al. Diet induces parallel changes to the gut microbiota and problem solving performance in a wild bird. Sci Rep 2020; 10: 20783.

8. Velando A, Noguera JC, Aira M, Domínguez J. Gut microbiome and telomere length in gull hatchlings. Biol Lett 2021; 17: 20210398.

9. Gensollen T, Iyer SS, Kasper DL, Blumberg RS. How colonization by microbiota in early life shapes the immune system. Science 2016; 352: 539–544.

10. Simon K, Verwoolde MB, Zhang J, Smidt H, de Vries Reilingh G, Kemp B, et al. Long-term effects of early life microbiota disturbance on adaptive immunity in laying hens. Poultry Science 2016; 95: 1543–1554.

11. Knutie SA, Wilkinson CL, Kohl KD, Rohr JR. Early-life disruption of amphibian microbiota decreases later-life resistance to parasites. Nat Commun 2017; 8: 86.

12. Kirschman LJ, Khadjinova A, Ireland K, Milligan-Myhre KC. Early life disruption of the microbiota affects organ development and cytokine gene expression in Threespine Stickleback. Integr Comp Biol 2020; icaa136.

13. Trevelline BK, MacLeod KJ, Knutie SA, Langkilde T, Kohl KD. *In ovo* microbial communities: a potential mechanism for the initial acquisition of gut microbiota among oviparous birds and lizards. Biol Lett 2018; 14: 20180225.

14. Teyssier A, Lens L, Matthysen E, White J. Dynamics of gut microbiota diversity during the early development of an avian host: Evidence from a cross-foster experiment. Front Microbiol 2018; 9: 1524.

15. Chen C-Y, Chen C-K, Chen Y-Y, Fang A, Shaw GT-W, Hung C-M, et al. Maternal gut microbes shape the early-life assembly of gut microbiota in passerine chicks via nests. Microbiome 2020; 8: 129.

16. Teyssier A, Matthysen E, Hudin NS, de Neve L, White J, Lens L. Diet contributes to urban-induced alterations in gut microbiota: experimental evidence from a wild passerine. Proc R Soc B 2020; 287: 20192182.

17. Dion-Phénix H, Charmantier A, de Franceschi C, Bourret G, Kembel SW, Réale D. Bacterial microbiota similarity between predators and prey in a blue tit trophic network. ISME J 2021; 15: 1098–1107.

18. Bodawatta KH, Freiberga I, Puzejova K, Sam K, Poulsen M, Jønsson KA. Flexibility and resilience of Great Tit (*Parus major*) gut microbiomes to changing diets. Anim microbiome 2021; 3: 20.

19. Hird SM, Carstens BC, Cardiff SW, Dittmann DL, Brumfield RT. Sampling locality is more detectable than taxonomy or ecology in the gut microbiota of the brood-parasitic Brown-headed Cowbird (*Molothrus ater*). PeerJ 2014; 2: e321.

20. Grond K, Santo Domingo JW, Lanctot RB, Jumpponen A, Bentzen RL, Boldenow ML, et al. Composition and drivers of gut microbial communities in arctic-breeding shorebirds. Front Microbiol 2019; 10: 2258.

21. Loo WT, García-Loor J, Dudaniec RY, Kleindorfer S, Cavanaugh CM. Host phylogeny, diet, and habitat differentiate the gut microbiomes of Darwin’s finches on Santa Cruz Island. Sci Rep 2019; 9: 18781.

22. Herder EA, Spence AR, Tingley MW, Hird SM. Elevation correlates with significant changes in relative abundance in hummingbird fecal microbiota, but composition changes little. Front Ecol Evol 2021; 8: 597756.

23. Drobniak SM, Cichoń M, Janas K, Barczyk J, Gustafsson L, Zagalska-Neubauer M. Habitat shapes diversity of gut microbiomes in a wild population of blue tits *Cyanistes caeruleus*. J Avian Biol 2021; jav.02829.

24. Starck JM, Ricklefs RE. Patterns of development: The Altricial-Precocial spectrum. In: Starck JM, Ricklefs RE (eds). Avian growth and development. Evolution within the altricial-precocial spectrum. 1998. Oxford University Press, New York, pp 2–30.

25. Grond K, Lanctot RB, Jumpponen A, Sandercock BK. Recruitment and establishment of the gut microbiome in arctic shorebirds. FEMS Microbiol Ecol 2017; 93: fix142.

26. Benskin CMcWH, Rhodes G, Pickup RW, Mainwaring MC, Wilson K, Hartley IR. Life history correlates of fecal bacterial species richness in a wild population of the blue tit *Cyanistes caeruleus*. Ecol Evol 2015; 5: 821–835.

27. Kreisinger J, Kropáčková L, Petrželková A, Adámková M, Tomášek O, Martin J-F, et al. Temporal stability and the effect of transgenerational transfer on fecal microbiota structure in a long distance migratory bird. Front Microbiol 2017; 8: 50.

28. Davidson GL, Somers SE, Wiley N, Johnson CN, Reichert MS, Ross RP, et al. A time-lagged association between the gut microbiome, nestling weight and nestling survival in wild Great Tits. J Anim Ecol 2021; 90: 989–1003.

29. Goodenough AE, Stallwood B, Dandy S, Nicholson TE, Stubbs H, Coker DG. Like mother like nest: similarity in microbial communities of adult female Pied Flycatchers and their nests. J Ornithol 2017; 158: 233–244.

30. Devaynes A, Antunes A, Bedford A, Ashton P. Progression in the bacterial load during the breeding season in nest boxes occupied by the Blue Tit and its potential impact on hatching or fledging success. J Ornithol 2018; 159: 1009–1017.

31. Moore PR, Evenson A, Luckey TD, McCoy E, Elvehjem CA, Hart EB. Use of sulfasuxidine, streptothricin, and streptomycin in nutritional studies with the chick. J Biol Chem 1946; 165: 437–441.

32. Jukes TH, Williams WL. Nutritional effects of antibiotics. Pharmacol Rev 1953; 5: 381–420.

33. Reuben RC, Roy PC, Sarkar SL, Alam R-U, Jahid IK. Isolation, characterization, and assessment of lactic acid bacteria toward their selection as poultry probiotics. BMC Microbiol 2019; 19: 253.

34. Dumonceaux TJ, Hill JE, Hemmingsen SM, Van Kessel AG. Characterization of intestinal microbiota and response to dietary Virginiamycin supplementation in the broiler chicken. Appl Environ Microbiol 2006; 72: 2815–2823.

35. Potti J, Moreno J, Yorio P, Briones V, García-Borboroglu P, Villar S, et al. Bacteria divert resources from growth for Magellanic Penguin chicks: Bacteria affect penguin chick growth. Ecol Lett 2002; 5: 709–714.

36. Kohl KD, Brun A, Bordenstein SR, Caviedes-Vidal E, Karasov WH. Gut microbes limit growth in house sparrow nestlings (*Passer domesticus*) but not through limitations in digestive capacity. Integr Zool 2018; 13: 139–151.

37. Knutie SA. Food supplementation affects gut microbiota and immunological resistance to parasites in a wild bird species. J Appl Ecol 2020; 57: 536–547.

38. Murray MH, Lankau EW, Kidd AD, Welch CN, Ellison T, Adams HC, et al. Gut microbiome shifts with urbanization and potentially facilitates a zoonotic pathogen in a wading bird. PLoS ONE 2020; 15: e0220926.

39. Berlow M, Phillips JN, Derryberry EP. Effects of urbanization and landscape on gut microbiomes in White-Crowned Sparrows. Microb Ecol 2021; 81: 253–266.

40. Berlow M, Wada H, Derryberry EP. Experimental exposure to noise alters gut microbiota in a captive songbird. Microb Ecol 2021; https://doi.org/10.1007/s00248-021-01924-3.

41. Lucas FS, Heeb P. Environmental factors shape cloacal bacterial assemblages in Great Tit *Parus major* and Blue Tit *P. caeruleus* nestlings. J Avian Biol 2005; 36: 510–516.

42. Perrins CM. British tits. 1979. HarperCollins, London.

43. Møller AP, Adriaensen F. Variation in clutch size in relation to nest size in birds. Ecol Evol 2014; 4: 3583–3595.

44. Gibb J. The breeding biology of the Great and Blue Titmice. Ibis 1950; 92: 507–539.

45. Perrins CM. Tits and their caterpillar food supply. Ibis 1991; 133: 49–54.

46. García-Navas V, Ferrer ES, Sanz JJ. Prey choice, provisioning behaviour, and effects of early nutrition on nestling phenotype of titmice. Écoscience 2013; 20: 9–18.

47. Barrientos R, Bueno-Enciso J, Sanz JJ. Hatching asynchrony vs. foraging efficiency: the response to food availability in specialist vs. generalist tit species. Sci Rep 2016; 6: 37750.

48. Bodawatta KH, Klečková I, Klečka J, Pužejová K, Koane B, Poulsen M, et al. Specific gut bacterial responses to natural diets of tropical birds. Sci Rep 2022; 12: 713.

49. Bodawatta KH, Puzejova K, Sam K, Poulsen M, Jønsson KA. Cloacal swabs and alcohol bird specimens are good proxies for compositional analyses of gut microbial communities of Great Tits (*Parus major*). Anim microbiome 2020; 2: 9.

50. Callahan BJ, McMurdie PJ, Rosen MJ, Han AW, Johnson AJA, Holmes SP. DADA2: High-resolution sample inference from Illumina amplicon data. Nat Methods 2016; 13: 581– 583.

51. Bolyen E, Rideout JR, Dillon MR, Bokulich NA, Abnet CC, Al-Ghalith GA, et al. Reproducible, interactive, scalable and extensible microbiome data science using QIIME 2. Nat Biotechnol 2019; 37: 852–857.

52. Quast C, Pruesse E, Yilmaz P, Gerken J, Schweer T, Yarza P, et al. The SILVA ribosomal RNA gene database project: improved data processing and web-based tools. Nucleic Acids Res 2012; 41: 590–596.

53. McMurdie PJ, Holmes S. phyloseq: An R package for reproducible interactive analysis and graphics of microbiome census data. PLoS ONE 2013; 8: e61217.

54. R Core Team. R: A language and environment for statistical computing. 2021. R Foundation for Statistical Computing, Viena, Austria.

55. Lahti L, Shetty S. Tools for microbiome analysis in R. 2017.

56. Kembel SW, Cowan PD, Helmus MR, Cornwell WK, Morlon H, Ackerly DD, et al. Picante: R tools for integrating phylogenies and ecology. Bioinformatics 2010; 26: 1463– 1464.

57. Bates D, Maechler M, Bolker BM, Walker S. Fitting linear mixed-effects models using lme4. J Stat Softw 2015; 67: 1–48.

58. Bartón K. MuMIn: multi-model inference. 2015. R package.

59. Nakagawa S, Schielzeth H. A general and simple method for obtaining R2 from generalized linear mixed-effects models. Methods Ecol Evol 2013; 4: 133–142.

60. Oksanen J, Blanchet FG, Friendly M, Kindt R, Legendre P, McGlinn D, et al. vegan: Community Ecology Package. 2020.

61. Arbizu PM. pairwiseAdonis: Pairwise multilevel comparison using adonis. 2018.

62. Liu C, Cui Y, Li X, Yao M. *microeco* : an R package for data mining in microbial community ecology. FEMS Microbiol Ecol 2021; 97: fiaa255.

63. Kurtz ZD, Müller CL, Miraldi ER, Littman DR, Blaser MJ, Bonneau RA. Sparse and Compositionally Robust Inference of Microbial Ecological Networks. PLoS Comput Biol 2015; 11: e1004226.

64. Csardi G, Nepusz T. The igraph software package for complex network research. Int J Complex Syst 2006; 1695: 1–9.

65. Bastian M, Heymann S, Jacomy M. Gephi: an open source software for exploring and manipulating networks. 2009.

66. Conway JR, Lex A, Gehlenborg N. UpSetR: an R package for the visualization of intersecting sets and their properties. Bioinformatics 2017; 33: 2938–2940.

67. Pebesma E. Simple Features for R: Standardized Support for Spatial Vector Data. The R Journal 2018; 10: 439.

68. Krams I, Vrublevska J, Cirule D, Kivleniece I, Krama T, Rantala MJ, et al. Stress, behaviour and immunity in wild caught wintering Great Tits (*Parus major*). Ethology 2013; 10.

69. Fischer CP, Wright-Lichter J, Romero LM. Chronic stress and the introduction to captivity: How wild house sparrows (*Passer domesticus*) adjust to laboratory conditions. Gen Comp Endocrinol 2018; 259: 85–92.

70. Bodawatta KH, Koane B, Maiah G, Sam K, Poulsen M, Jønsson KA. Species-specific but not phylosymbiotic gut microbiomes of New Guinean 2 passerines are shaped by diet and flight-associated gut modifications. Proc R Soc Lond B 2021; 288: 20210446.

71. Hinde CA, Kilner RM. Negotiations within the family over the supply of parental care. Proc R Soc B 2007; 274: 53–60.

72. Wilkin TA, King LE, Sheldon BC. Habitat quality, nestling diet, and provisioning behaviour in great tits *Parus major*. J Avian Biol 2009; 40: 135–145.

73. Tremblay I, Thomas D, Blondel J, Perret P, Lambrechts MM. The effect of habitat quality on foraging patterns, provisioning rate and nestling growth in Corsican Blue Tits *Parus caeruleus*. Ibis 2004; 147: 17–24.

74. Dickens M, Berridge D, Hartley IR. Biparental care and offspring begging strategies: hungry nestling blue tits move towards the father. Anim Behav 2008; 75: 167–174.

75. Santema P, Schlicht E, Kempenaers B. Testing the conditional cooperation model: what can we learn from parents taking turns when feeding offspring? Front Ecol Evol 2019; 7: 94.

76. Mainwaring MC. Causes and consequences of intraspecific variation in nesting behaviors: Insights from Blue Tits and Great Tits. Front Ecol Evol 2017; 5: 39.

77. Pendlebury CJ, Bryant DM. Night-time behaviour of egg-laying tits. Ibis 2005; 147: 342–345.

78. Lord AM, Mccleery RH, Cresswell W. Incubation prior to clutch completion accelerates embryonic development and so hatch date for eggs laid earlier in a clutch in the Great Tit *Parus major*. J Avian Biol 2011; 42: 187–191.

79. Diez-Méndez D, Sanz JJ, Barba E. Impacts of ambient temperature and clutch size on incubation behaviour onset in a female-only incubator songbird. Ibis 2021; 163: 1056–1071.

80. Nilsson J-Å. Time-dependent reproductive decisions in the blue tit. Oikos 2000; 88: 351–361.

81. Bambini G, Schlicht E, Kempenaers B. Patterns of female nest attendance and male feeding throughout the incubation period in Blue Tits *Cyanistes caeruleus*. Ibis 2019; 161: 50–65.

82. Diez-Méndez D, Cooper CB, Sanz JJ, Verdejo J, Barba E. Deconstructing incubation behaviour in response to ambient temperature over different timescales. J Avian Biol 2021; jav.02781.

83. Rodríguez S, Barba E. Nestling growth is impaired by heat stress: an experimental study in a mediterranean Great Tit population. Zool Stud 2016; 55: 40.

84. Andreasson F, Nord A, Nilsson J-Å. Brood size constrains the development of endothermy in blue tits. J Exp Biol 2016; 219: 2212–2219.

85. Hird SM, Ganz H, Eisen JA, Boyce WM. The cloacal microbiome of five wild duck species varies by species and influenza A virus infection status. mSphere 2018; 3: e00382–18.

86. Constable GWA, Fagan B, Law R. Maternal transmission as a symbiont sieve, and the absence of lactation in male mammals. 2022. https://doi.org/10.1101/2022.01.10.475639.

87. Frank SA. Host–symbiont conflict over the mixing of symbiotic lineages. Proc R Soc Lond B 1996; 263: 339–344.

88. Naef-Daenzer B. Patch time allocation and patch sampling by foraging great and blue tits. Anim Behav 2000; 59: 989–999.

89. Minot EO, Perrins CM. Interspecific interference competition - nest sites for blue and great tits. J Anim Ecol 1986; 55: 331–350.

90. Kempenaers B, Dhondt AA. Competition between Blue and Great tit for roosting sites in winter: an aviary experiment. Ornis Scand 1991; 22: 73–75.

91. Stauss MJ, Burkhardt JF, Tomiuk J. Foraging flight distances as a measure of parental effort in blue tits *Parus caeruleus* differ with environmental conditions. J Avian Biol 2005; 36: 47–56.

92. Bańbura J, Lambrechts MM, Blondel J, Perret P, Cartan-Son M. Food handling time of blue tits chicks: constraints and adaptation to different prey types. J Avian Biol 1999; 30: 263–270.

