## Supplementary Material for "Gut microbiome disturbances of altricial Blue and Great tit nestlings are countered by continuous microbial inoculations from parental microbiomes"

**Table S1**: Great tit data from the efficacy experiment showing the number of the metallic ring, the sex, and the treatment of an individual. Weight data shows the change in body mass since the day they were mist-netted (day 0) until the day they were released back to their habitat (day 8).

| | | | **Weight (g)** | | | | | |
|---|---|---|---|---|---|---|---|---|
| **Ring no.** | **Sex** | **Treatment** | **Day 0** | **Day 1** | **Day 2** | **Day 3** | **Day 4** | **Day 8** |
| N941680 | Male | Antibiotics | 18.32 | 16.54 | 16.54 | 16.09 | 16.28 | 15.48 |
| N941682 | Male | Probiotics | 18.48 | 17.10 | 16.69 | 16.61 | 16.40 | 15.94 |
| N941677 | Male | Antibiotics | 19.38 | 17.39 | 17.45 | 16.72 | 16.75 | 15.97 |
| N941678 | Female | Probiotics | 17.78 | 16.15 | 16.00 | 15.86 | 15.51 | 14.95 |
| N941676 | Female | Antibiotics | 17.15 | 15.80 | 15.41 | 15.35 | 15.52 | 14.30 |
| N941698 | Male | Probiotics | 19.03 | 18.19 | 17.25 | 17.05 | 16.56 | 15.72 |
| N941699 | Male | Antibiotics | 17.87 | 17.53 | 16.51 | 16.30 | 16.33 | 16.39 |
| N941697 | Male | Probiotics | 17.53 | 16.37 | 15.87 | 15.74 | 15.55 | 15.70 |
| N930804 | Male | Antibiotics | 17.50 | 17.60 | 17.27 | 17.15 | 17.07 | 16.59 |
| N930805 | Male | Probiotics | 16.99 | 16.93 | 16.73 | 16.60 | 16.87 | 16.16 |

**Table S2**: Doses applied to the experimental hatchlings during the field experiment. Antibiotic and probiotic doses were diluted in a water volume similar to the indicated for control doses.

| | Day 1 | Day 4 | Day 7 | Day 10 | Day 13 |
|---|---|---|---|---|---|
| **Great tits** | | | | | |
| Expected mean weight per chick (g) | 1 | 5 | 10 | 15 | 18 |
| Antibiotic (mg) | 0.5 | 2.5 | 5.0 | 7.5 | 9.0 |
| Probiotic (mg) | 6.7 | 33.5 | 67.0 | 100.5 | 120.6 |
| Control (ml) | 0.1 | 0.15 | 0.175 | 0.2 | 0.2 |
| **Blue tits** | | | | | |
| Expected mean weight per chick (g) | 1 | 3 | 6 | 9 | 11 |
| Antibiotic (mg) | 0.5 | 1.5 | 3.0 | 4.5 | 5.5 |
| Probiotic (mg) | 6.7 | 20.1 | 40.2 | 60.3 | 73.7 |
| Control (ml) | 0.1 | 0.15 | 0.175 | 0.2 | 0.2 |

**Table S3**. Estimates resulting from the linear mixed-effect models of antibiotic and probiotic treatments, day of experiment and host sex on alpha diversities of gut microbiomes in the efficacy experiment. The categorical variable *treatment* was assessed in reference to the probiotic treatment. Individual ID was a random factor. Significant results are indicated in bold font.

| | Estimate | SE | t value | p |
|---|---|---|---|---|
| **Observed ASV richness** | | | | |
| $R^2_{fixed} = 0.29$, $R^2_{fixed+random\ effects} = 0.63$ | | | | |
| Intercept | 246.90 | 86.965 | 2.839 | |
| Treatment | -5.18 | 63.072 | -0.082 | 0.937 |
| **Day** | **-485.06** | **181.48** | **-2.673** | **0.011** |
| $Day^2$ | 86.68 | 140.495 | 0.617 | 0.541 |
| Weight | -1.60 | 35.375 | -0.045 | 0.964 |
| Sex | -57.56 | 94.743 | -0.608 | 0.556 |
| Treatment*Day | -12.62 | 186.789 | -0.068 | 0.947 |
| Treatment*$Day^2$ | 190.49 | 189.013 | 1.008 | 0.321 |
| **Faith's PD** | | | | |
| $R^2_{fixed} = 0.27$, $R^2_{fixed+random\ effects} = 0.46$ | | | | |
| Intercept | 11.14 | 1.882 | 5.922 | |
| Treatment | 0.09 | 1.290 | 0.069 | 0.947 |
| **Day** | **-13.97** | **4.941** | **-2.827** | **0.007** |
| $Day^2$ | 5.30 | 4.027 | 1.315 | 0.197 |
| Weight | -0.38 | 0.904 | -0.419 | 0.678 |
| Sex | -0.56 | 2.097 | -0.265 | 0.796 |
| Treatment*Day | 2.97 | 5.391 | 0.551 | 0.586 |
| Treatment*$Day^2$ | 0.30 | 5.444 | 0.055 | 0.957 |
| **Shannon's diversity index** | | | | |
| $R^2_{fixed} = 0.38$, $R^2_{fixed+random\ effects} = 0.60$ | | | | |
| Intercept | 4.86 | 0.709 | 6.861 | |
| Treatment | -0.79 | 0.496 | -1.589 | 0.156 |
| Day | -2.31 | 1.714 | -1.346 | 0.186 |
| $Day^2$ | -0.78 | 1.369 | -0.573 | 0.571 |
| Weight | 0.48 | 0.322 | 1.48 | 0.149 |
| Sex | -1.03 | 0.784 | -1.319 | 0.213 |
| Treatment*Day | -1.48 | 1.827 | -0.812 | 0.423 |
| Treatment*$Day^2$ | 1.33 | 1.847 | 0.722 | 0.476 |
| **Relative dominance** (log transformed) | | | | |
| $R^2_{fixed} = 0.38$, $R^2_{fixed+random\ effects} = 0.60$ | | | | |
| Intercept | -2.52 | 0.445 | -5.673 | |
| Treatment | 0.63 | 0.307 | 2.048 | 0.080 |
| Day | 0.65 | 1.132 | 0.579 | 0.566 |
| $Day^2$ | 1.10 | 0.916 | 1.198 | 0.239 |
| Weight | -0.39 | 0.210 | -1.853 | 0.074 |
| Sex | 0.57 | 0.494 | 1.145 | 0.275 |
| Treatment*Day | 0.57 | 1.225 | 0.462 | 0.647 |
| Treatment*$Day^2$ | -0.60 | 1.237 | -0.486 | 0.630 |

**Table S4:** Estimates resulting from the linear mixed-effect models of control, antibiotic and probiotic treatments and day of experiment on alfa diversities of gut microbiomes in wild chicks (field experiment). The categorical variable *treatment* was assessed in reference to the control treatment. Brood ID was a random intercept factor and chick ID was a random slope factor within day of experiment. Significant results are indicated with bold font. Table is attached as a separate excel file.

| Great tits | Estimate | SE | t value | p |
|---|---|---|---|---|
| **Observed ASV richness** (square root transf.) | | | | |
| $R^2_{fixed} = 0.40$, $R^2_{fixed+random\ effects} = 0.50$ | | | | |
| Intercept | 13.24 | 0.574 | 23.074 | |
| **Antibiotic** | 1.01 | 0.548 | 1.841 | **0.067** |
| Probiotic | 0.55 | 0.552 | 0.99 | 0.323 |
| **Day** | **44.25** | **7.362** | **6.011** | **<0.001** |
| **Day$^2$** | **-35.48** | **6.838** | **-5.188** | **<0.001** |
| Antibiotic*Day | 3.67 | 10.309 | 0.356 | 0.723 |
| Antibiotic *Day$^2$ | -3.22 | 10.309 | -0.312 | 0.756 |
| Probiotic*Day | -2.81 | 9.547 | -0.295 | 0.769 |
| Probiotic *Day$^2$ | 4.26 | 9.550 | 0.446 | 0.656 |
| **Faith's PD** | | | | |
| $R^2_{fixed} = 0.32$, $R^2_{fixed+random\ effects} = 0.39$ | | | | |
| Intercept | 10.49 | 0.427 | 24.576 | |
| **Antibiotic** | **0.86** | **0.429** | **2.005** | **0.046** |
| Probiotic | 0.52 | 0.433 | 1.211 | 0.227 |
| **Day** | **29.46** | **5.292** | **5.567** | **<0.001** |
| **Day$^2$** | **-24.15** | **5.239** | **-4.61** | **<0.001** |
| Antibiotic*Day | -0.38 | 7.425 | -0.051 | 0.959 |
| Antibiotic *Day$^2$ | -7.02 | 7.417 | -0.947 | 0.345 |
| Probiotic*Day | -3.07 | 7.322 | -0.419 | 0.675 |
| Probiotic *Day$^2$ | 4.29 | 7.336 | 0.584 | 0.560 |
| **Shannon's diversity index** (log transf.) | | | | |
| $R^2_{fixed} = 0.38$, $R^2_{fixed+random\ effects} = 0.60$ | | | | |
| Intercept | 18.58 | 0.966 | 19.245 | |
| Antibiotic | -0.40 | 0.803 | -0.497 | 0.620 |
| Probiotic | 0.04 | 0.804 | 0.045 | 0.965 |
| **Day** | **44.48** | **10.636** | **4.182** | **<0.001** |
| **Day$^2$** | **-58.66** | **11.138** | **-5.266** | **<0.001** |
| Antibiotic*Day | 2.29 | 15.153 | 0.151 | 0.880 |
| Antibiotic *Day$^2$ | -7.25 | 14.978 | -0.484 | 0.630 |
| Probiotic*Day | 1.25 | 15.928 | 0.078 | 0.938 |
| Probiotic *Day$^2$ | 8.72 | 15.785 | 0.553 | 0.584 |
| **Relative dominance** (log transf.) | | | | |
| $R^2_{fixed} = 0.38$, $R^2_{fixed+random\ effects} = 0.60$ | | | | |
| Intercept | -2.33 | 0.090 | -25.967 | |
| Antibiotic | 0.06 | 0.08 | 0.749 | 0.455 |
| Probiotic | -0.00 | 0.083 | -0.022 | 0.982 |
| Day | -2.02 | 1.204 | -1.677 | 0.099 |
| **Day$^2$** | **4.11** | **1.211** | **3.393** | **0.001** |
| Antibiotic*Day | 1.45 | 1.699 | 0.856 | 0.396 |
| Antibiotic *Day$^2$ | 1.17 | 1.704 | 0.689 | 0.494 |
| Probiotic*Day | -0.33 | 1.701 | -0.196 | 0.845 |
| Probiotic *Day$^2$ | -0.61 | 1.710 | -0.357 | 0.722 |

| Blue tits | Estimate | SE | t value | p |
|---|---|---|---|---|
| **Observed ASV richness** (square root transf.) | | | | |
| $R^2_{fixed} = 0.41$, $R^2_{fixed+random\ effects} = 0.61$ | | | | |
| Intercept | 12.83 | 0.641 | 20.020 | |
| Antibiotic | -0.32 | 0.713 | -0.447 | 0.658 |
| Probiotic | -0.77 | 0.720 | -1.063 | 0.295 |
| **Day** | **38.25** | **7.224** | **5.295** | **<0.001** |
| **Day$^2$** | **-20.88** | **6.222** | **-3.357** | **0.001** |
| Antibiotic*Day | 6.053 | 9.826 | 0.616 | 0.542 |
| Antibiotic *Day$^2$ | 6.012 | 10.270 | 0.585 | 0.562 |
| Probiotic*Day | -2.83 | 8.552 | -0.331 | 0.742 |
| Probiotic *Day$^2$ | -8.77 | 8.831 | -0.993 | 0.323 |
| **Faith's PD** | | | | |
| $R^2_{fixed} = 0.32$, $R^2_{fixed+random\ effects} = 0.39$ | | | | |
| Intercept | 10.98 | 0.495 | 22.198 | |
| Antibiotic | -0.40 | 0.581 | -0.696 | 0.490 |
| Probiotic | -0.70 | 0.587 | -1.184 | 0.244 |
| **Day** | **29.30** | **5.310** | **5.519** | **<0.001** |
| **Day$^2$** | **-14.00** | **5.032** | **-2.783** | **0.006** |
| Antibiotic*Day | 2.08 | 7.246 | 0.288 | 0.775 |
| Antibiotic *Day$^2$ | -1.14 | 7.581 | -0.150 | 0.881 |
| Probiotic*Day | -6.61 | 6.899 | -0.958 | 0.340 |
| Probiotic *Day$^2$ | -3.56 | 7.149 | -0.498 | 0.619 |
| **Shannon's diversity index** | | | | |
| $R^2_{fixed} = 0.32$, $R^2_{fixed+random\ effects} = 0.61$ | | | | |
| Intercept | 4.16 | 0.140 | 29.589 | |
| Antibiotic | -0.08 | 0.138 | -0.612 | 0.545 |
| Probiotic | -0.17 | 0.138 | -1.238 | 0.225 |
| **Day** | **6.54** | **1.442** | **4.533** | **<0.001** |
| **Day$^2$** | **-2.88** | **1.219** | **-2.361** | **0.021** |
| Antibiotic*Day | 0.98 | 1.961 | 0.499 | 0.621 |
| Antibiotic *Day$^2$ | 0.44 | 2.047 | 0.213 | 0.833 |
| Probiotic*Day | -0.76 | 1.675 | -0.454 | 0.651 |
| Probiotic *Day$^2$ | -1.97 | 1.726 | -1.143 | 0.257 |
| **Relative dominance** (log transf.) | | | | |
| $R^2_{fixed} = 0.14$, $R^2_{fixed+random\ effects} = 0.60$ | | | | |
| Intercept | -2.33 | 0.134 | -17.387 | |
| Antibiotic | 0.06 | 0.108 | 0.583 | 0.564 |
| Probiotic | 0.02 | 0.102 | 0.197 | 0.845 |
| Day | -1.95 | 1.340 | -1.453 | 0.156 |
| Day$^2$ | 0.86 | 0.895 | 0.965 | 0.337 |
| Antibiotic*Day | -3.01 | 1.812 | -1.661 | 0.107 |
| Antibiotic *Day$^2$ | -1.95 | 1.895 | -1.029 | 0.311 |
| Probiotic*Day | 0.12 | 1.226 | 0.099 | 0.921 |
| Probiotic *Day$^2$ | 0.36 | 1.276 | 0.284 | 0.777 |

**Table S5**: Nest-box coordinates in decimal degrees where experimental Great tits (GT) and Blue tits (BT) bred.

| Nest-box ID | Species | Latitude | Longitude |
|---|---|---|---|
| 6 | BT | 48.975208 | 14.427872 |
| 7 | GT | 48.975376 | 14.427562 |
| 13 | GT | 48.97437 | 14.425271 |
| 14 | BT | 48.973945 | 14.425397 |
| 18 | BT | 48.973658 | 14.424246 |
| 20 | GT | 48.973796 | 14.42289 |
| 23 | GT | 48.973734 | 14.421232 |
| 29 | GT | 48.972894 | 14.416866 |
| 32 | GT | 48.9739 | 14.415584 |
| 37 | GT | 48.975645 | 14.415549 |
| 36 | GT | 48.976257 | 14.416205 |
| 34 | BT | 48.976951 | 14.416727 |
| 74 | GT | 48.978074 | 14.417141 |
| 78 | GT | 48.978969 | 14.415103 |
| 70 | BT | 48.97913 | 14.414627 |
| 90 | BT | 48.984384 | 14.416846 |
| C15 | BT | 48.98453 | 14.4207 |
| 68 | BT | 48.984724 | 14.421852 |
| 93 | BT | 48.984421 | 14.423256 |
| 100 | BT | 48.983528 | 14.426514 |

**Table S6**: Estimates resulting from the linear mixed-effect models of control, antibiotic and probiotic treatments and day of experiment on body mass gain in wild chicks. Brood ID is random intercept factor. Chick ID is a random slope intercept factor within day of experiment. The categorical variable *treatment* was assessed in reference to the control treatment. Significant results are indicated with bold font.

| Great tits | Estimate | SE | t value | p |
|---|---|---|---|---|
| $R^2_{fixed} = 0.97$, $R^2_{fixed+random\ effects} = 0.98$ | | | | |
| Intercept | 9.07 | 0.140 | 64.992 | |
| Antibiotic | 0.08 | 0.173 | 0.449 | 0.656 |
| Probiotic | 0.18 | 0.175 | 1.015 | 0.316 |
| **Day** | **95.71** | **1.392** | **68.732** | **<0.001** |
| **Day$^2$** | **-12.72** | **1.416** | **-8.982** | **<0.001** |
| Antibiotic*Day | -2.15 | 1.971 | -1.089 | 0.279 |
| Antibiotic *Day$^2$ | -0.15 | 1.992 | -0.074 | 0.941 |
| Probiotic*Day | -1.97 | 2.014 | -0.977 | 0.331 |
| Probiotic*Day$^2$ | -0.37 | 2.036 | -0.182 | 0.856 |
| Blue tits | | | | |
| $R^2_{fixed} = 0.91$, $R^2_{fixed+random\ effects} = 0.98$ | | | | |
| Intercept | 4.78 | 0.197 | 24.304 | |
| Antibiotic | 0.07 | 0.270 | 0.255 | 0.800 |
| Probiotic | 0.19 | 0.272 | 0.698 | 0.489 |
| **Day** | **48.01** | **1.998** | **24.03** | **<0.001** |
| Day$^2$ | -2.93 | 1.762 | -1.662 | 0.103 |
| Antibiotic*Day | 0.78 | 2.725 | 0.287 | 0.775 |
| Antibiotic *Day$^2$ | 1.14 | 2.813 | 0.405 | 0.688 |
| Probiotic*Day | -0.72 | 2.411 | -0.299 | 0.766 |
| Probiotic*Day$^2$ | -2.43 | 2.467 | -0.984 | 0.329 |

**Table S7**: Estimates resulting from the linear mixed-effect models of control, antibiotic and probiotic treatments and untreated chicks on body size (tarsus length) at day 16. Brood ID is random intercept factor. The categorical variable *treatment* was assessed in reference to the control treatment. Significant results are indicated with bold font.

| Great tits | Estimate | SE | t value | p |
|---|---|---|---|---|
| $R^2_{fixed} = 0.06$, $R^2_{fixed+random\ effects} = 0.23$ | | | | |
| Intercept | 19.88 | 0.114 | 174.967 | |
| Controls | 0.23 | 0.138 | 1.685 | 0.097 |
| Antibiotics | 0.01 | 0.138 | 0.037 | 0.970 |
| Probiotics | 0.27 | 0.140 | 1.933 | 0.057 |

| Blue tits | | | | |
|---|---|---|---|---|
| $R^2_{fixed} = 0.01$, $R^2_{fixed+random\ effects} = 0.39$ | | | | |
| Intercept | 16.55 | 0.200 | 82.558 | |
| Controls | 0.09 | 0.229 | 0.390 | 0.699 |
| Antibiotics | -0.13 | 0.209 | -0.613 | 0.543 |
| Probiotics | -0.02 | 0.244 | -0.073 | 0.942 |

**Table S8**: Estimates resulting from the linear mixed-effect models of control, antibiotic and probiotic treatments and untreated chicks, together with nest and parents on alpha diversities at day 16. Brood ID was a random intercept factor. The categorical variable *treatment* was assessed in reference to the control treatment. Significant results are indicated with bold font. Table is attached as a separate excel file.

| Great tits | Estimate | SE | t value | p |
|---|---|---|---|---|
| **Observed ASV richness** | | | | |
| $R^2_{fixed} = 0.16$, $R^2_{fixed+random\ effects} = 0.32$ | | | | |
| Intercept | 247.82 | 25.485 | 9.724 | |
| Antibiotic | 33.64 | 30.276 | 1.111 | 0.270 |
| Probiotic | 11.78 | 30.698 | 0.384 | 0.702 |
| Untreated | 4.62 | 28.015 | 0.165 | 0.869 |
| **Nest** | **103.38** | **35.801** | **2.888** | **0.005** |
| Female | 78.97 | 51.234 | 1.541 | 0.127 |
| **Male** | **-75.24** | **34.772** | **-2.164** | **0.033** |
| **Faith's PD** | | | | |
| $R^2_{fixed} = 0.05$, $R^2_{fixed+random\ effects} = 0.16$ | | | | |
| Intercept | 11.85 | 0.631 | 18.786 | |
| Antibiotic | 0.58 | 0.805 | 0.717 | 0.475 |
| Probiotic | 0.16 | 0.816 | 0.191 | 0.849 |
| Untreated | 0.05 | 0.745 | 0.066 | 0.947 |
| Nest | 0.99 | 0.953 | 1.041 | 0.301 |
| Female | 2.15 | 1.359 | 1.579 | 0.118 |
| Male | -0.68 | 0.925 | -0.737 | 0.463 |
| **Shannon's diversity index** | | | | |
| $R^2_{fixed} = 0.16$, $R^2_{fixed+random\ effects} = 0.35$ | | | | |
| Intercept | 4.28 | 0.223 | 19.169 | |
| Antibiotic | 0.05 | 0.258 | 0.190 | 0.850 |
| Probiotic | 0.08 | 0.261 | 0.306 | 0.760 |
| Untreated | -0.07 | 0.239 | -0.285 | 0.776 |
| Nest | 0.53 | 0.305 | 1.733 | 0.087 |
| Female | -0.46 | 0.436 | -1.056 | 0.294 |
| **Male** | **-1.05** | **0.296** | **-3.541** | **<0.001** |

| Blue tits | Estimate | SE | t value | p |
|---|---|---|---|---|
| **Observed ASV richness** | | | | |
| $R^2_{fixed} = 0.28$, $R^2_{fixed+random\ effects} = 0.32$ | | | | |
| Intercept | 274.26 | 42.528 | 6.449 | |
| Antibiotic | -10.31 | 53.460 | -0.193 | 0.848 |
| Probiotic | -28.35 | 58.901 | -0.481 | 0.632 |
| Untreated | -9.05 | 48.709 | -0.186 | 0.853 |
| **Nest** | **210.50** | **58.282** | **3.612** | **<0.001** |
| Female | -56.54 | 58.651 | -0.964 | 0.339 |
| Male | -26.93 | 63.546 | -0.424 | 0.673 |
| **Faith's PD** | | | | |
| $R^2_{fixed} = 0.16$, $R^2_{fixed+random\ effects} = 0.29$ | | | | |
| Intercept | 13.82 | 1.090 | 12.673 | |
| Antibiotic | -1.15 | 1.306 | -0.883 | 0.381 |
| Probiotic | -1.71 | 1.446 | -1.183 | 0.242 |
| Untreated | -0.73 | 1.191 | -0.615 | 0.541 |
| Nest | 2.75 | 1.418 | 1.937 | 0.058 |

| | | | | |
|---|---|---|---|---|
| Female | -1.85 | 1.439 | -1.284 | 0.204 |
| Male | -1.52 | 1.561 | -0.975 | 0.334 |
| **Shannon's diversity index** | | | | |
| $R^2_{fixed} = 0.29$, $R^2_{fixed+random\ effects} = 0.44$ | | | | |
| Intercept | 4.80 | 0.295 | 16.292 | |
| Antibiotic | -0.13 | 0.342 | -0.369 | 0.713 |
| Probiotic | -0.41 | 0.379 | -1.090 | 0.280 |
| Untreated | -0.08 | 0.312 | -0.244 | 0.808 |
| Nest | 0.40 | 0.371 | 1.078 | 0.286 |
| **Female** | **-1.35** | **0.378** | **-3.574** | **<0.001** |
| **Male** | **-1.20** | **0.410** | **-2.915** | **0.005** |

**Table S9:** Pairwise comparisons of chick, adult and nest microbiome composition on final day based on Bray-Curtis distances. Significant results are indicated with bold font.

| | F statistic | $R^2$ | Adjusted p |
|---|---|---|---|
| Great tits | | | |
| **Male vs. Control chicks** | **4.003** | **0.1291** | **0.002** |
| **Male vs. Antibiotic chicks** | **4.129** | **0.1371** | **0.002** |
| **Male vs. Probiotic chicks** | **4.097** | **0.1408** | **0.002** |
| **Male vs. Untreated chicks** | **4.536** | **0.1209** | **0.002** |
| Male vs. Female | 0.809 | 0.0632 | 0.999 |
| **Male vs. Nest** | **3.353** | **0.1571** | **0.002** |
| Female vs. Control chicks | 1.843 | 0.0807 | 0.067 |
| Female vs. Antibiotic chicks | 2.002 | 0.0911 | 0.050 |
| Female vs. Probiotic chicks | 2.025 | 0.0963 | 0.069 |
| **Female vs. Untreated chicks** | **2.066** | **0.0023** | **0.048** |
| Female vs. Nest | 1.774 | 0.1288 | 0.092 |
| Control vs. Antibiotic chicks | 0.533 | 0.0151 | 0.999 |
| Control vs. Probiotic chicks | 0.449 | 0.0131 | 0.999 |
| Control vs. Untreated chicks | 0.621 | 0.0146 | 0.999 |
| Control chicks vs. Nest | 0.847 | 0.0304 | 0.999 |
| Antibiotic vs. Probiotic chicks | 0.441 | 0.0132 | 0.999 |
| Antibiotic vs. Untreated chicks | 0.649 | 0.0156 | 0.999 |
| Antibiotic vs. Nest | 0.884 | 0.0329 | 0.999 |
| Probiotic vs. Untreated chicks | 0.585 | 0.0144 | 0.999 |
| Probiotic chicks vs. Nest | 0.873 | 0.0338 | 0.999 |
| Untreated chicks vs. Nest | 1.271 | 0.0371 | 0.999 |
| Blue tits | | | |
| **Male vs. Control chicks** | **2.123** | **0.1503** | **0.025** |
| **Male vs. Antibiotic chicks** | **2.334** | **0.1273** | **0.004** |
| **Male vs. Probiotic chicks** | **2.081** | **0.1478** | **0.023** |
| **Male vs. Untreated chicks** | **2.445** | **0.0891** | **0.019** |
| Male vs. Female | 0.799 | 0.0624 | 0.999 |
| **Male vs. Nest** | **2.359** | **0.1643** | **0.011** |
| Female vs. Control chicks | 1.934 | 0.1214 | 0.265 |
| Female vs. Antibiotic chicks | 2.035 | 0.1016 | 0.050 |
| Female vs. Probiotic chicks | 1.702 | 0.1084 | 0.905 |
| **Female vs. Untreated chicks** | **2.175** | **0.0745** | **0.036** |
| Female vs. Nest | 2.117 | 0.1313 | 0.130 |
| Control vs. Antibiotic chicks | 0.847 | 0.0449 | 0.999 |
| Control vs. Probiotic chicks | 0.872 | 0.0586 | 0.999 |
| Control vs. Untreated chicks | 0.763 | 0.0275 | 0.999 |
| Control chicks vs. Nest | 0.997 | 0.0665 | 0.999 |
| Antibiotic vs. Probiotic chicks | 0.838 | 0.0445 | 0.999 |
| Antibiotic vs. Untreated chicks | 0.721 | 0.0227 | 0.999 |
| Antibiotic vs. Nest | 0.994 | 0.0523 | 0.999 |
| Probiotic vs. Untreated chicks | 0.768 | 0.0277 | 0.999 |
| Probiotic chicks vs. Nest | 0.961 | 0.0643 | 0.999 |
| Untreated chicks vs. Nest | 1.017 | 0.0363 | 0.999 |

**Table S10:** Estimates resulting from the linear mixed-effect models of nest and chick categories and day of experiment on alpha diversities. Brood ID is random intercept factor. The categorical variable *chicks* was assessed in reference to *nests*. Significant results are indicated with bold font.

| Great tits | Estimate | SE | t value | p |
|---|---|---|---|---|
| **Observed ASV richness** | | | | |
| $R^2_{fixed} = 0.48$, $R^2_{fixed+random\ effects} = 0.52$ | | | | |
| Intercept | 304.70 | 30.36 | 10.038 | |
| Day of experiment | 46.50 | 40.92 | 1.136 | 0.258 |
| **Chicks** | **-223.86** | **32.26** | **-6.939** | **<0.001** |
| **Day of experiment*Chicks** | **131.55** | **44.55** | **2.953** | **0.004** |
| **Faith's PD** | | | | |
| $R^2_{fixed} = 0.43$, $R^2_{fixed+random\ effects} = 0.48$ | | | | |
| Intercept | 12.26 | 0.785 | 15.625 | |
| Day of experiment | 0.58 | 1.067 | 0.541 | 0.589 |
| **Chicks** | **-4.85** | **0.841** | **-5.764** | **<0.001** |
| **Day of experiment*Chicks** | **4.01** | **1.161** | **3.456** | **<0.001** |
| **Shannon's diversity index** | | | | |
| $R^2_{fixed} = 0.29$, $R^2_{fixed+random\ effects} = 0.43$ | | | | |
| Intercept | 4.44 | 0.236 | 18.763 | |
| Day of experiment | 0.37 | 0.298 | 1.248 | 0.215 |
| **Chicks** | **-1.08** | **0.235** | **-4.585** | **<0.001** |
| Day of experiment*Chicks | 0.55 | 0.325 | 1.686 | 0.094 |

| Blue tits | Estimate | SE | t value | p |
|---|---|---|---|---|
| **Observed ASV richness** | | | | |
| $R^2_{fixed} = 0.63$, $R^2_{fixed+random\ effects} = 0.66$ | | | | |
| Intercept | 379.70 | 31.55 | 12.033 | |
| **Day of experiment** | **101.91** | **45.56** | **2.237** | **0.028** |
| **Chicks** | **-309.60** | **33.70** | **-9.187** | **<0.001** |
| Day of experiment*Chicks | 86.05 | 49.71 | 1.731 | 0.087 |
| **Faith's PD** | | | | |
| $R^2_{fixed} = 0.61$, $R^2_{fixed+random\ effects} = 0.65$ | | | | |
| Intercept | 13.74 | 0.787 | 17.455 | |
| **Day of experiment** | **2.58** | **1.121** | **2.304** | **0.023** |
| **Chicks** | **-6.47** | **0.829** | **-7.805** | **<0.001** |
| **Day of experiment*Chicks** | **2.84** | **1.223** | **2.323** | **0.022** |
| **Shannon's diversity index** | | | | |
| $R^2_{fixed} = 0.58$, $R^2_{fixed+random\ effects} = 0.60$ | | | | |
| Intercept | 4.99 | 0.193 | 25.852 | |
| Day of experiment | 0.15 | 0.282 | 0.538 | 0.592 |
| **Chicks** | **-1.67** | **0.209** | **-7.995** | **<0.001** |
| **Day of experiment*Chicks** | **1.16** | **0.308** | **3.751** | **<0.001** |

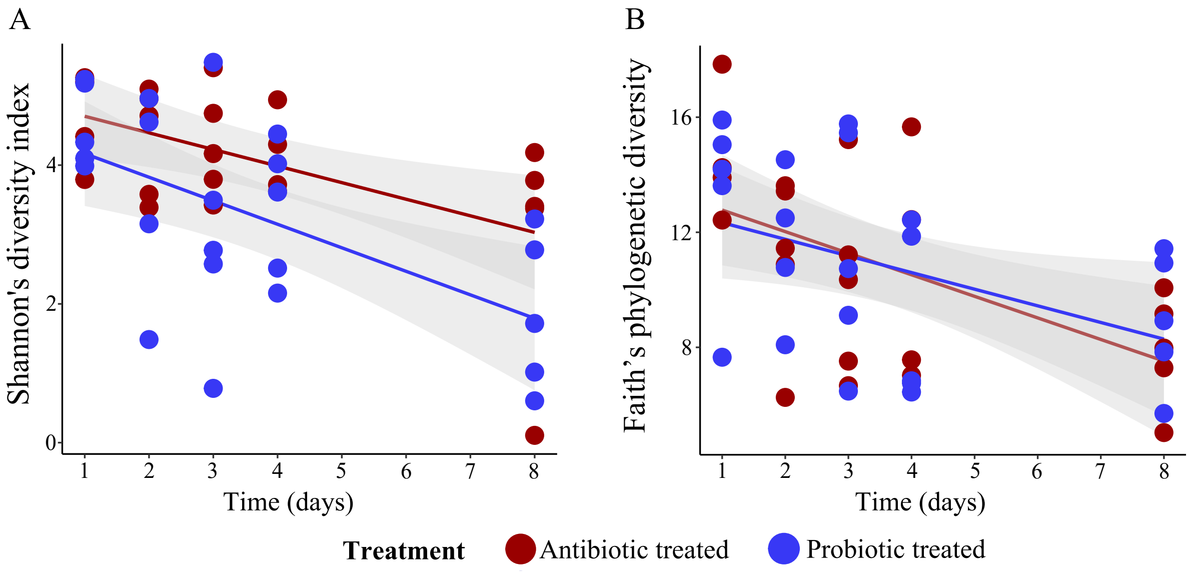

**Fig S1.** Alfa diversity patterns during the efficacy experiment: Shannon's diversity (A) and Faith's PD (B) in gut microbiomes of antibiotic and probiotic treated individuals.

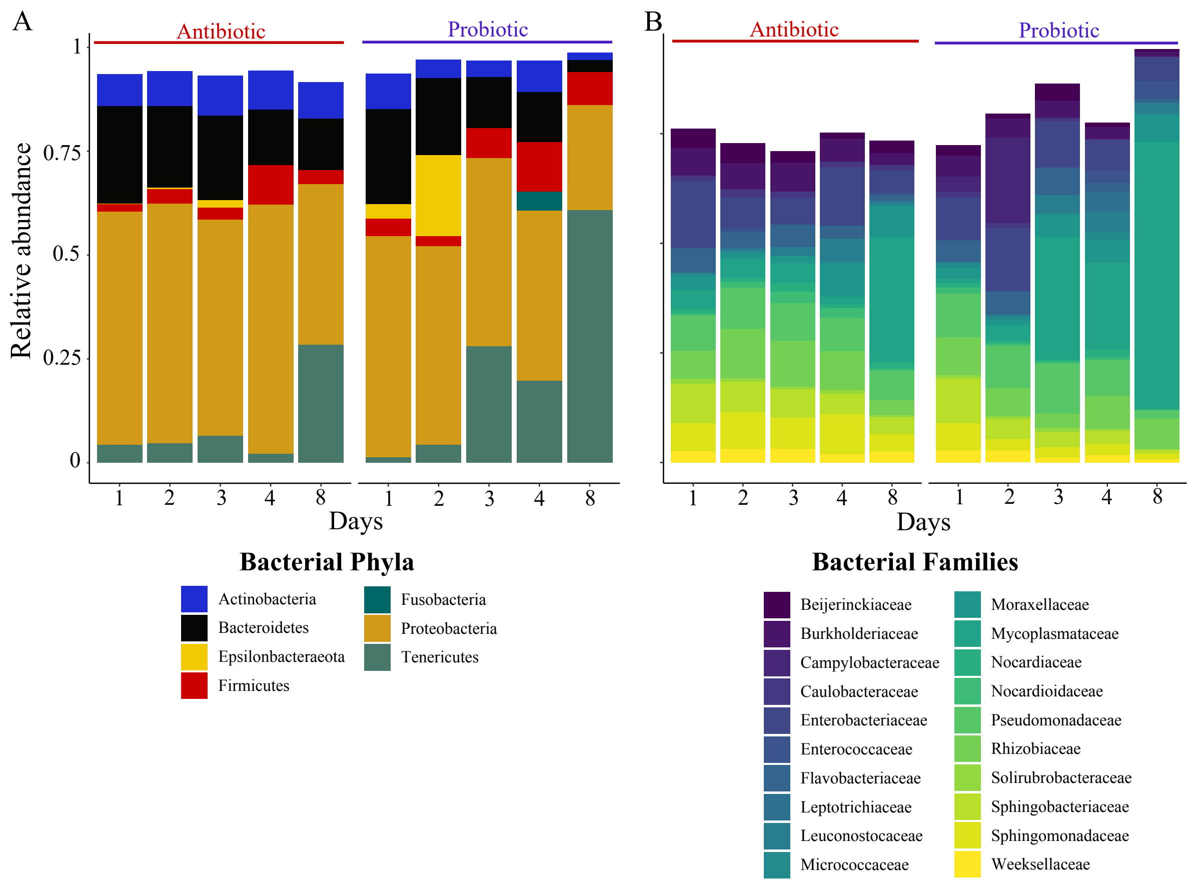

**Fig S2**. Relative abundance of main bacterial phyla (A) and the 20 most abundant bacterial families (B) in microbiomes of antibiotic and probiotic treated individuals across sampling days. Low abundant bacterial phyla and families are not shown in the figure.

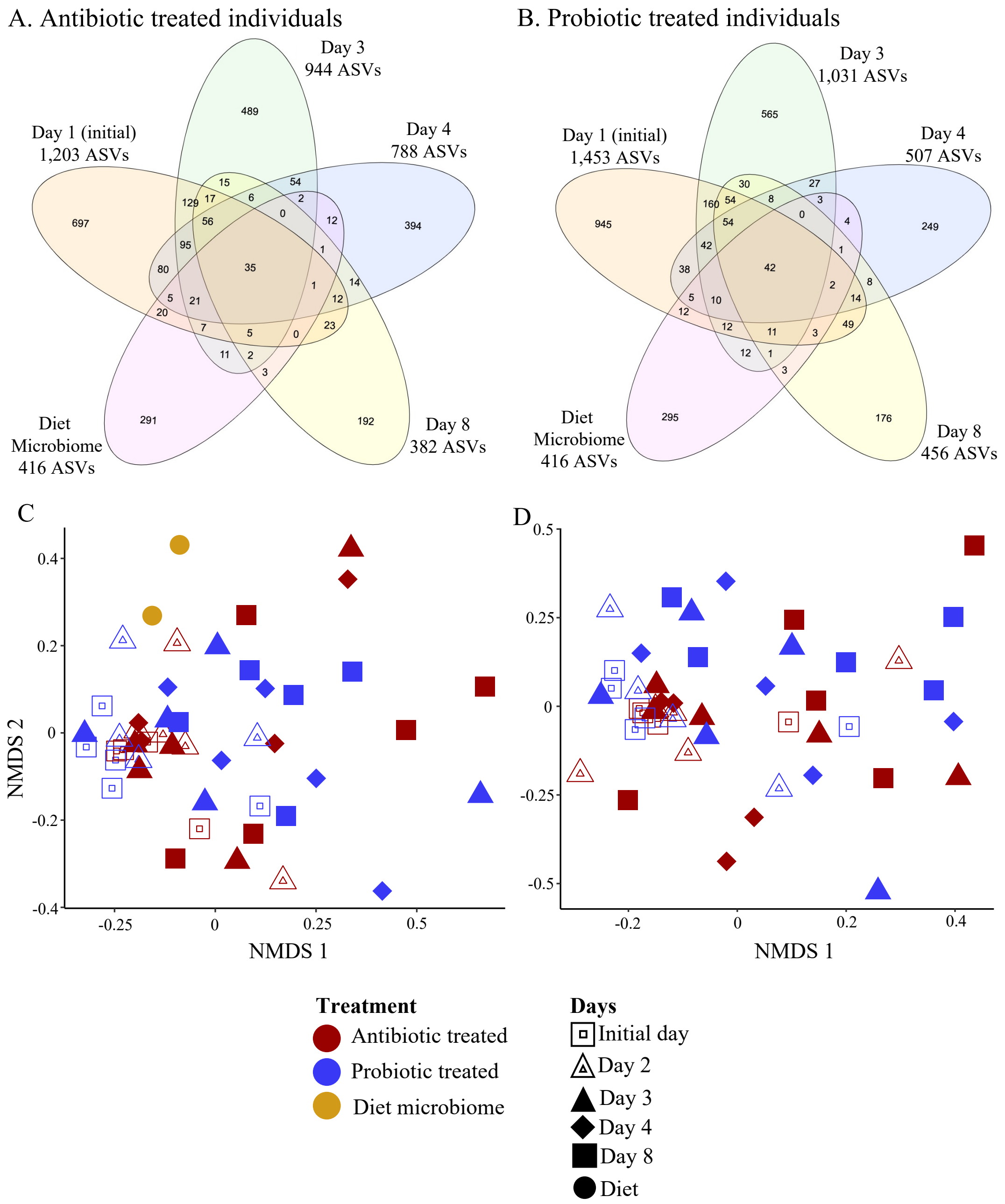

**Fig S3.** Flower plots depicting the shared ASVs in microbiomes of individuals during the treatment period (**A**: antibiotic treated, **B**: probiotic treated) and food microbiome in the efficacy experiment. NMDS plots showing the compositional similarity (based on Bray-Curtis distances) of gut microbiomes of treated individuals and the food microbiomes (stress = 0.2519) (**C**) and the compositional similarity of gut microbiomes after removing any ASV found in the food microbiome (stress = 0.2304) (**D**). The removal of food-born microbes from the analyses did not influence the statistical outcome of the effect of antibiotic/probiotic treatment over time (PERMANOVAs: Antibiotic $_{(Bray\text{-}Curtis)}$: $F = 1.063$, $R^2 = 0.1911$, $p = 0.2069$; Probiotic $_{(Bray\text{-}Curtis)}$: $F = 1.209$, $R^2 = 0.2029$, $p = 0.0997$).

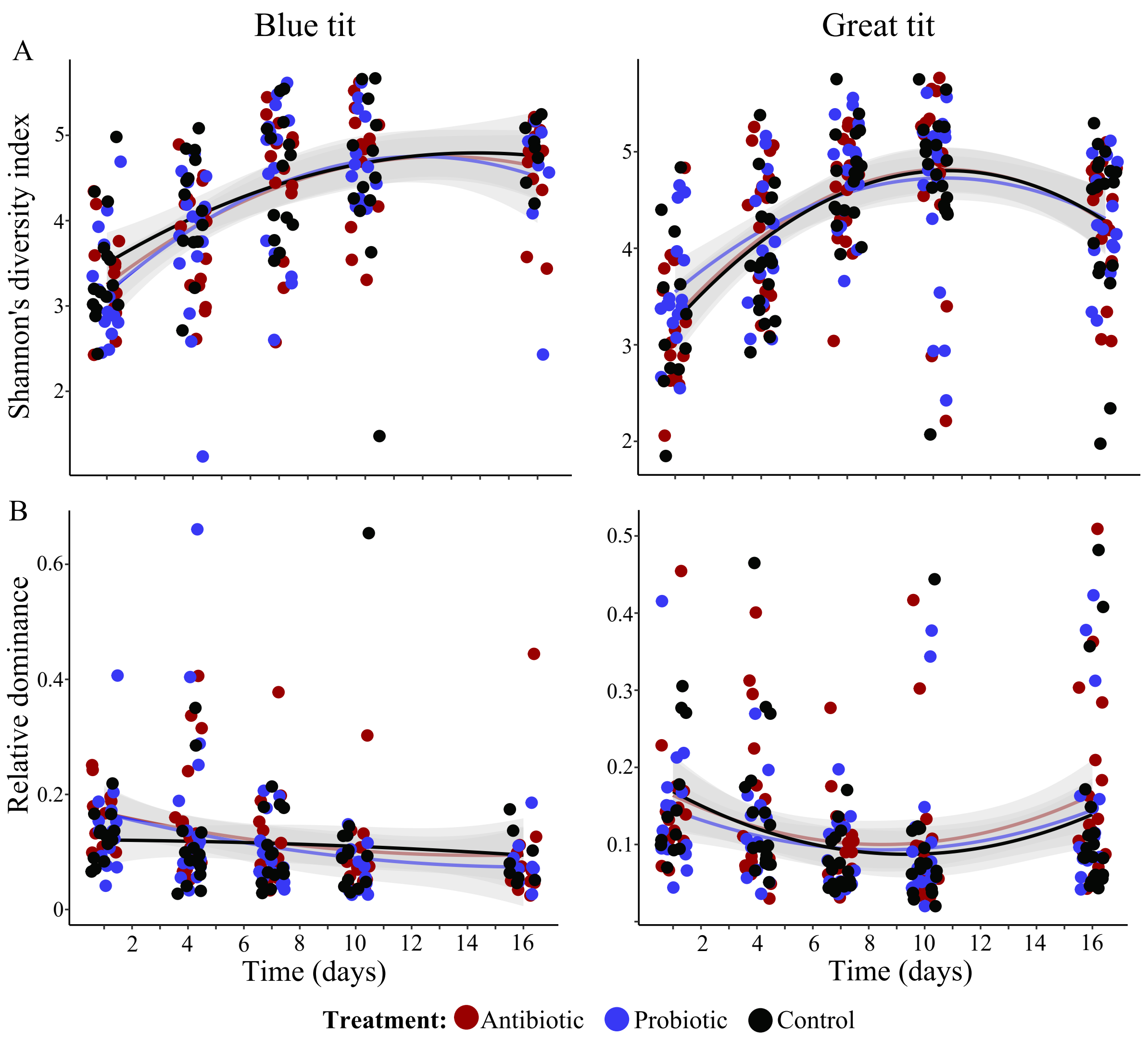

**Fig S4.** Association between days and Shannon's diversity index (A), and relative dominance (B) of gut microbial communities of developing chicks of *P. major* and *C. caeruleus*.

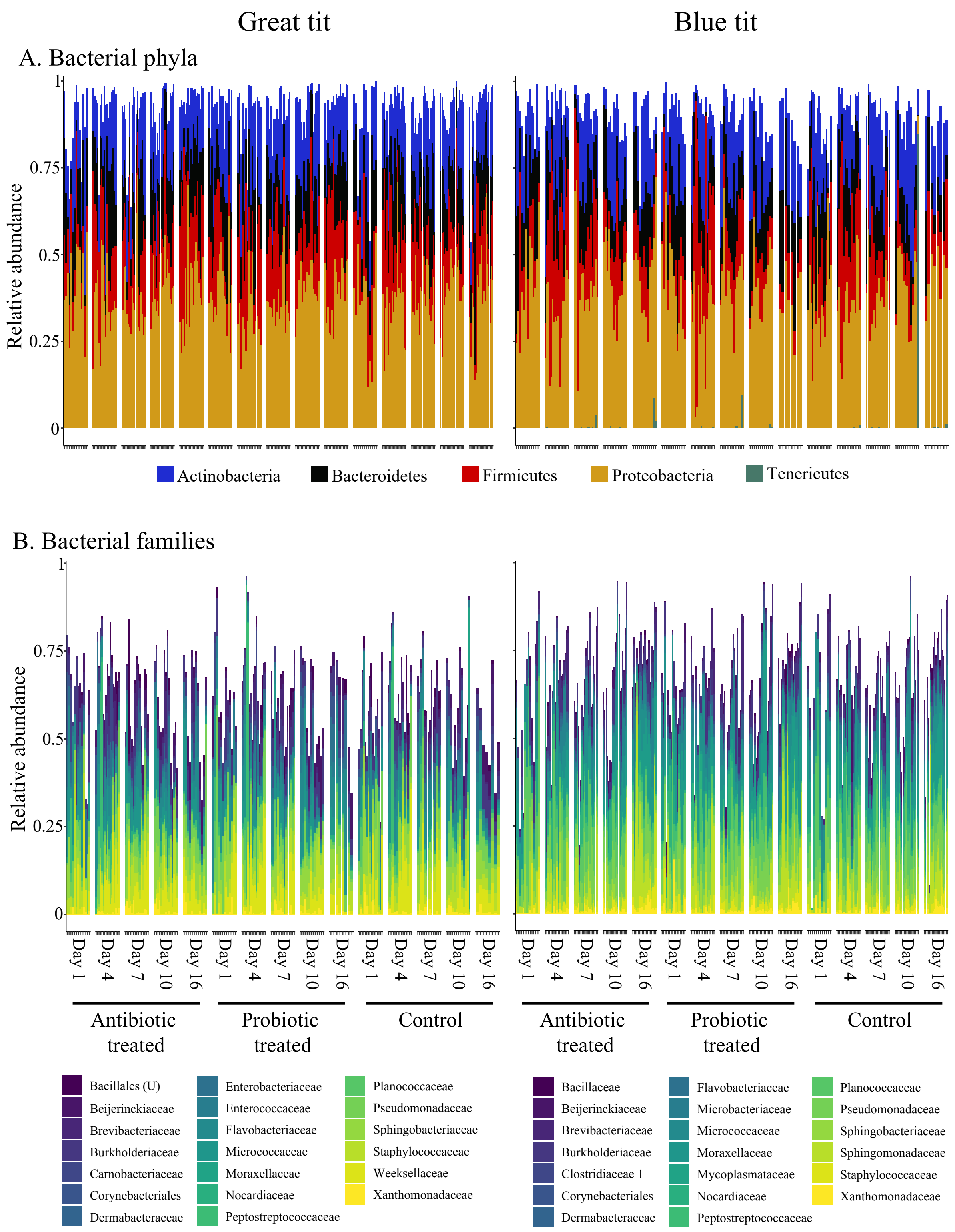

**Fig S5**. Relative abundance of main bacterial phyla (A) and 20 most abundant bacterial families (B) in microbiomes of chicks during the manipulation period. Each bar represent an individual and empty areas above each bar represent the relative abundance of other bacterial phyla and families.

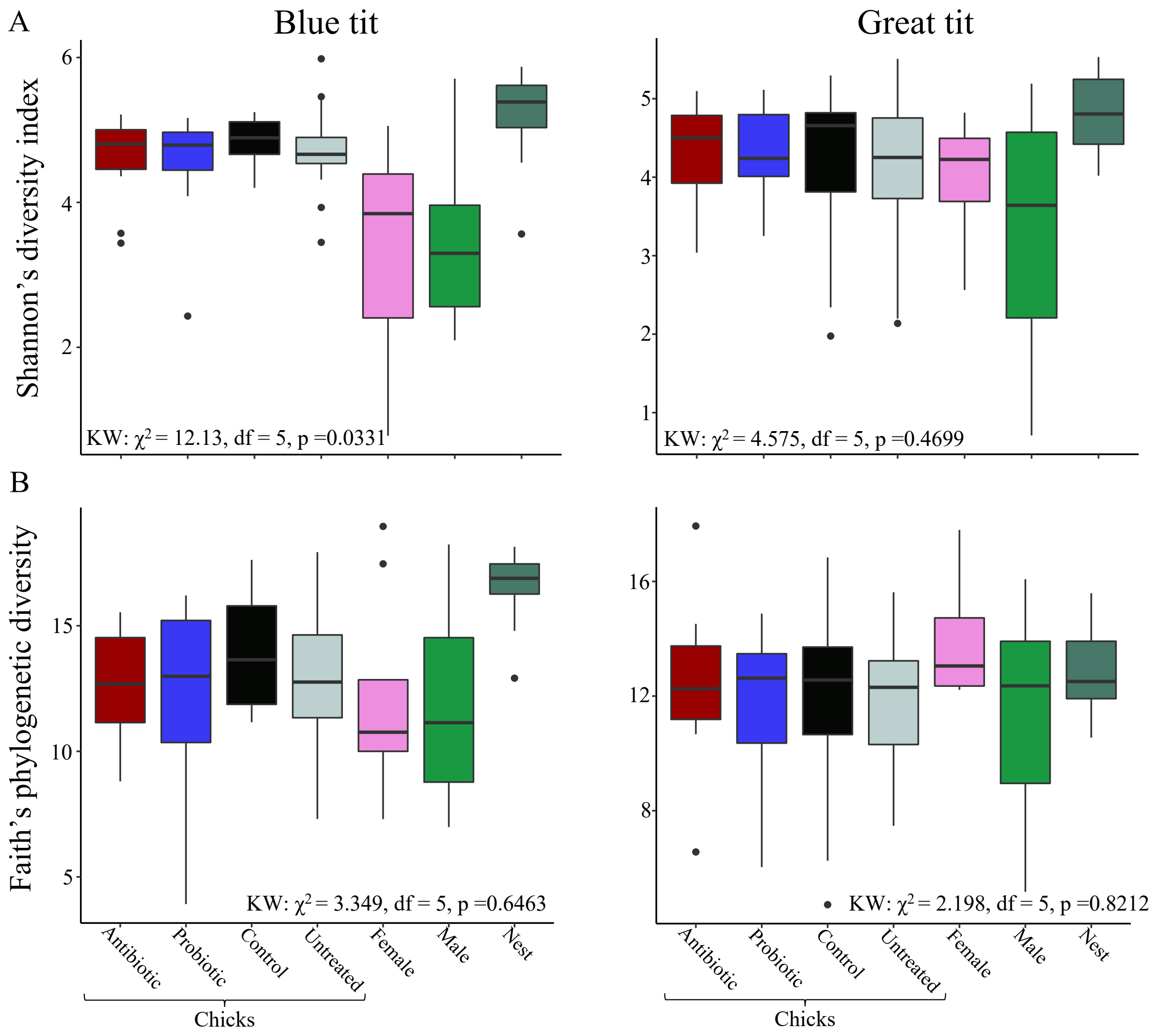

**Fig. S6**. Box plots depicting Shannon's diversity index (A) and Faith's phylogenetic diversity (B) of gut microbiomes of chicks and adults in the last sampling day.

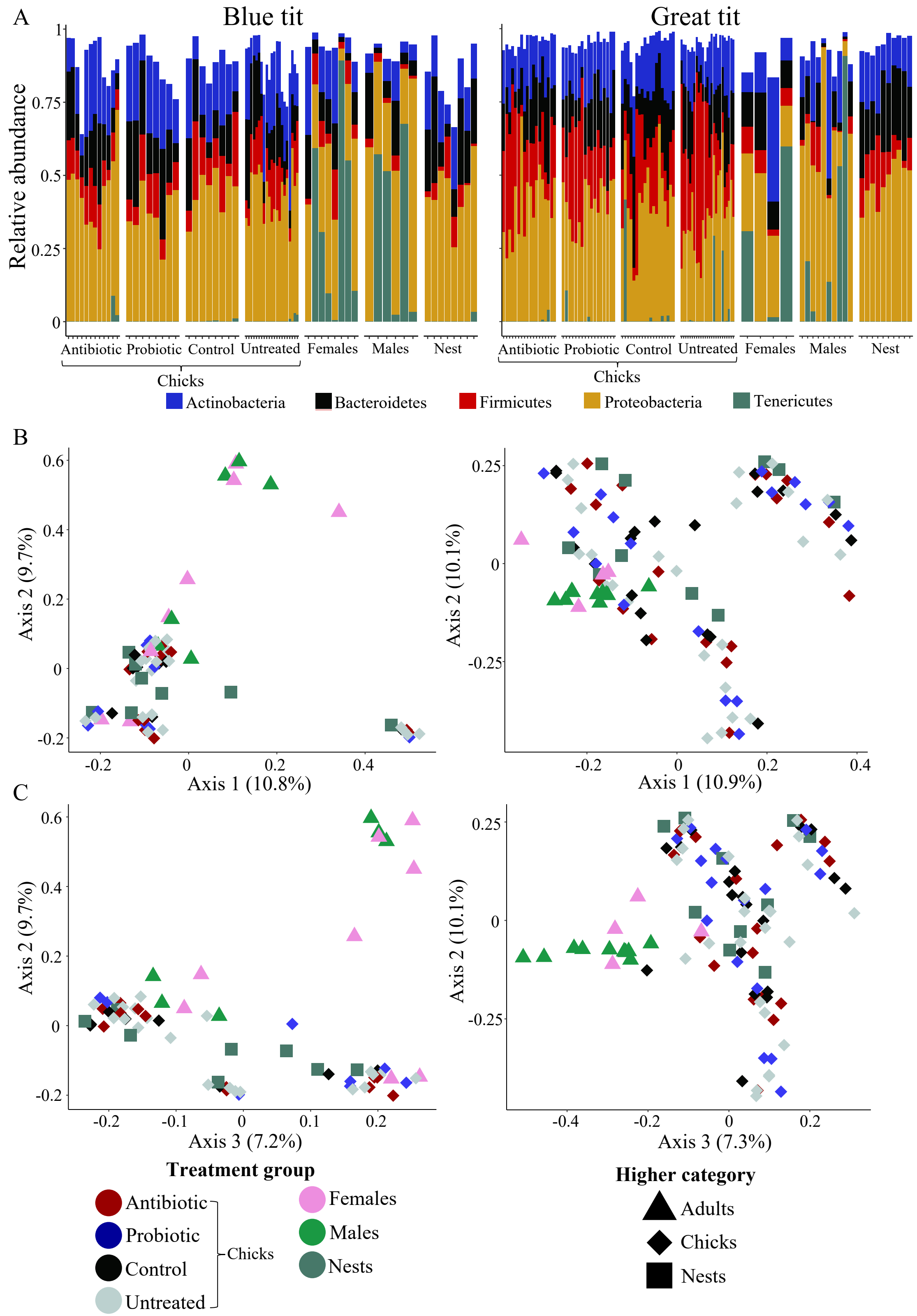

**Fig. S7**. (A) Relative abundance of dominant bacterial phyla in microbiomes of different treatment groups during the last sampling day. (B) Principal Coordinate Analyses (PCoA) plots (A: axis 1 and 2; B: Axis 2 and 3) depicting the microbial community similarity (Bray-Curtis distances) of different treatment groups during the last sampling day.

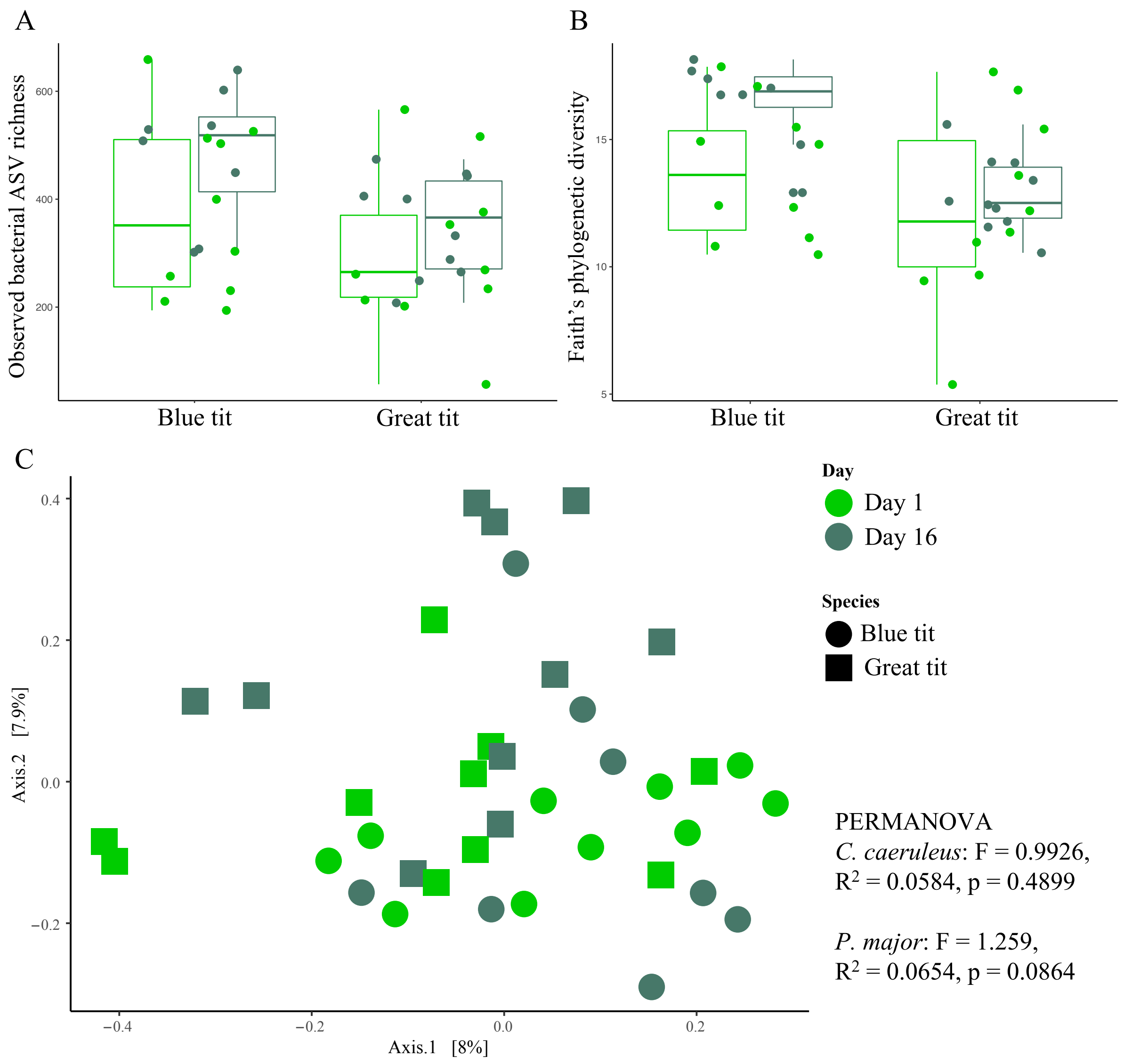

**Fig S8**. Observed ASV richness (A) and Faiths phylogenetic diversity (B) of nest microbiomes during 1st and last sampling days. (C) Nest bacterial community composition (measured with Bray-Curtis distances) during the two sampling time points.
